# A Novel *XACT* lncRNA Transcript with Functions Transcending X-Chromosome Inactivation

**DOI:** 10.1101/2025.01.03.631201

**Authors:** Peifen Zhang, Paula Haro Angles, Hong Zhang, Tingyue Li, Roger Mulet-Lazaro, Wilfred F.J. van IJcken, Sjaak Philipsen, Ruud Delwel, Joost Gribnau, Danny Huylebroeck, Frank Grosveld, Claire Rougeulle, Eskeatnaf Mulugeta

## Abstract

X-chromosome inactivation (XCI) is initiated during early mammalian embryogenesis by a long non-coding RNA (lncRNA) *XIST*, which coats one of the two X-chromosomes and facilitates epigenetic transcriptional silencing. A second, evolutionarily recent primate-specific lncRNA *XACT* was proposed to antagonize *XIST*’s ability to induce XCI. *XACT* expression is restricted to pluripotent states and early embryonic stages and coats the active X-chromosome in both females and males. Here, we report a novel *XACT* transcript expressed in normal and cancerous somatic cells from both the inactive (Xi) and active (Xa) X chromosomes. It coexists with *XIST* on the Xi without affecting *XIST* expression or (re)activating X-linked genes inactivated by XCI. During hematopoietic stem cell (HSC) differentiation, *XACT* is primarily expressed in myeloid progenitors in both sexes. *XACT* expression is activated in HSCs and peaks in megakaryocyte-erythrocyte progenitor cells (MEPs) before rapidly declining as the MEPs differentiate into megakaryocytes or erythrocytes. By combining CRISPR-based *XACT* perturbation with epigenomic and transcriptional studies, we revealed the key role of *XACT* in the self-renewal and differentiation of erythroid progenitors into erythrocytes, by recruiting cis-regulatory proteins and regulating transcription through ETS and AP-1 transcription factors. Furthermore, *XACT* is expressed in a subset of acute myeloid leukemia (AML) patients, with high levels found in erythroid-megakaryocytic blast cells; suggesting *XACT*’s potential as a marker for AML subtypes. Thus, we identified a novel *XACT* transcript with a unique expression profile and important roles in normal and cancer cells, revealing *XACT* lncRNA functions beyond its previously proposed role in XCI during embryogenesis.

## Introduction

Long non-coding RNAs (lncRNAs) are important transcripts that play critical roles in gene regulation and various cellular processes during healthy embryogenesis, cellular differentiation, and disease progression ^1-3^. However, the identities and mechanism(s) of action of lncRNAs remain largely unknown. One process that provided key insights into lncRNA function is X-chromosome inactivation (XCI) ^4,5^. In mammals, XCI compensates for gene-dosage inequality between males (XY) and females (XX), resulting from the evolutionary loss of most Y-chromosomal genes, by inactivating one of the two X-chromosomes in female somatic cells ^6-10^. XCI is triggered in early embryogenesis by the lncRNA *XIST*, which coats the chromosome and facilitates its epigenetic transcriptional silencing ^5,11-16^. In addition to *XIST*, other lncRNAs (including *Jpx*, *Tsix*, and *Xite*) have been identified as important for XCI ^11,17,18^. However, the exact mechanisms, partner regulatory proteins, and even the complete set of lncRNAs involved in XCI remain unclear, and differ among mammalian species ^19-21^. In recent years another lncRNA, *XACT* (X-active coating transcript), has been proposed to be a repressor of *XIST*’s ability to induce XCI ^22,23^.

*XACT* is a recent evolutionary addition specific to hominoids that emerged about 20 million years ago ^24^. Initially identified in human embryonic stem cells (hESCs), *XACT* is expressed from and coats the active X-chromosome (Xa) in both males and females ^22^. *XACT* encodes an exceptionally long, polyadenylated lncRNA, proposed to span 251.8 kilobases (kb), and is primarily localized in the nucleus^22^. In human pre-implantation embryos, *XACT* expression strongly correlates with that of *XIST* in early developmental stages ^22,23^. Its expression in pre-implantation embryos and hESCs is subsequently downregulated upon differentiation ^22,23^. *XACT* has also been implicated in the "erosion" of XCI in human pluripotent stem cells (hPSCs), characterized by a loss of XIST, reduction of repressive epigenetic signatures, and partial reactivation of X-linked genes ^23,25-27^. Despite these observations, the precise role of *XACT* in initiation or erosion of XCI is disputed. Follow-up investigations revealed that deletion of *XACT* in primed female hPSCs did not affect X-linked gene-dosage or allelic expression status, and the initiation of erosion depended more on intact *XIST* than on *XACT* ^28^. However, *XACT* deletion affected neural differentiation of primed hESCs ^28^. Recently, another deletion of *XACT* in naive hPSCs was shown neither to alter the expression of *XIST* nor global X-chromosome gene expression ^29,23^. Since its discovery, *XACT* expression has only been observed during early development, and its absence in differentiated cells shifted the focus towards its role in pluripotency and XCI in early development.

Here, we identified a hitherto unknown *XACT* transcript that is expressed in differentiated (male and female) normal and cancerous somatic cells, which is distinct from the transcript that is found in hESCs. The novel XACT transcript is expressed from both the active (Xa) and inactive (Xi) chromosomes and coexists with XIST on the Xi. However, *XACT* expression neither downregulates *XIST* nor reactivates genes silenced by XCI. This novel *XACT* transcript, discovered in K562 erythroleukemia cells, is also expressed in a subset of patients with acute myeloid leukemia (AML), in both sexes. Furthermore, during the differentiation of hematopoietic stem cells (HSCs), *XACT* is specifically expressed in megakaryocyte-erythrocyte progenitor cells (MEPs) of both sexes and is not found in other lineages. It is quickly downregulated during further MEP differentiation. Using HUDEP2 cells (an MEP-like cell line with erythroid differentiation potential), we show the key role of *XACT* in recruiting *cis*-regulatory proteins, regulating transcription, and maintaining the erythroid progenitor cell state, but also regulating the progenitors’ differentiation to erythrocytes through ETS and AP-1 family transcription factors (TFs). Finally, we also show that *XACT* can be used as a marker to discriminate subsets of AML patients. These results collectively reveal novel functions of *XACT* beyond XCI during embryogenesis.

## Results

### The *XACT* lncRNA is expressed in the K562 erythroleukemic cell line and AML patients

While optimizing a combinatorial barcoding-based single-cell RNA-sequencing (scRNA-seq) experiment (i.e. SPLiT-seq) ^30^ with pooled K562 (human, female) and MEL (mouse, male) erythroleukemia cell lines, we identified several lncRNAs among top 5 most abundant genes (Supplementary Table 1), including *XACT* transcripts in K562 cells. The visualization of the read alignments at the *XACT* locus revealed high expression of a previously undescribed transcript across a 443 kb-long region (Figure 1A). This new *XACT* transcript (chrX:113,616,300-114,059,121; GRCh38) is 191 kb longer than the originally described *XACT* transcript ^22,23,27^ (chrX:113,740,044-113,991,913, GRCh38) (Figure 1A, Supplementary Figure 1A).

**Figure 1.**
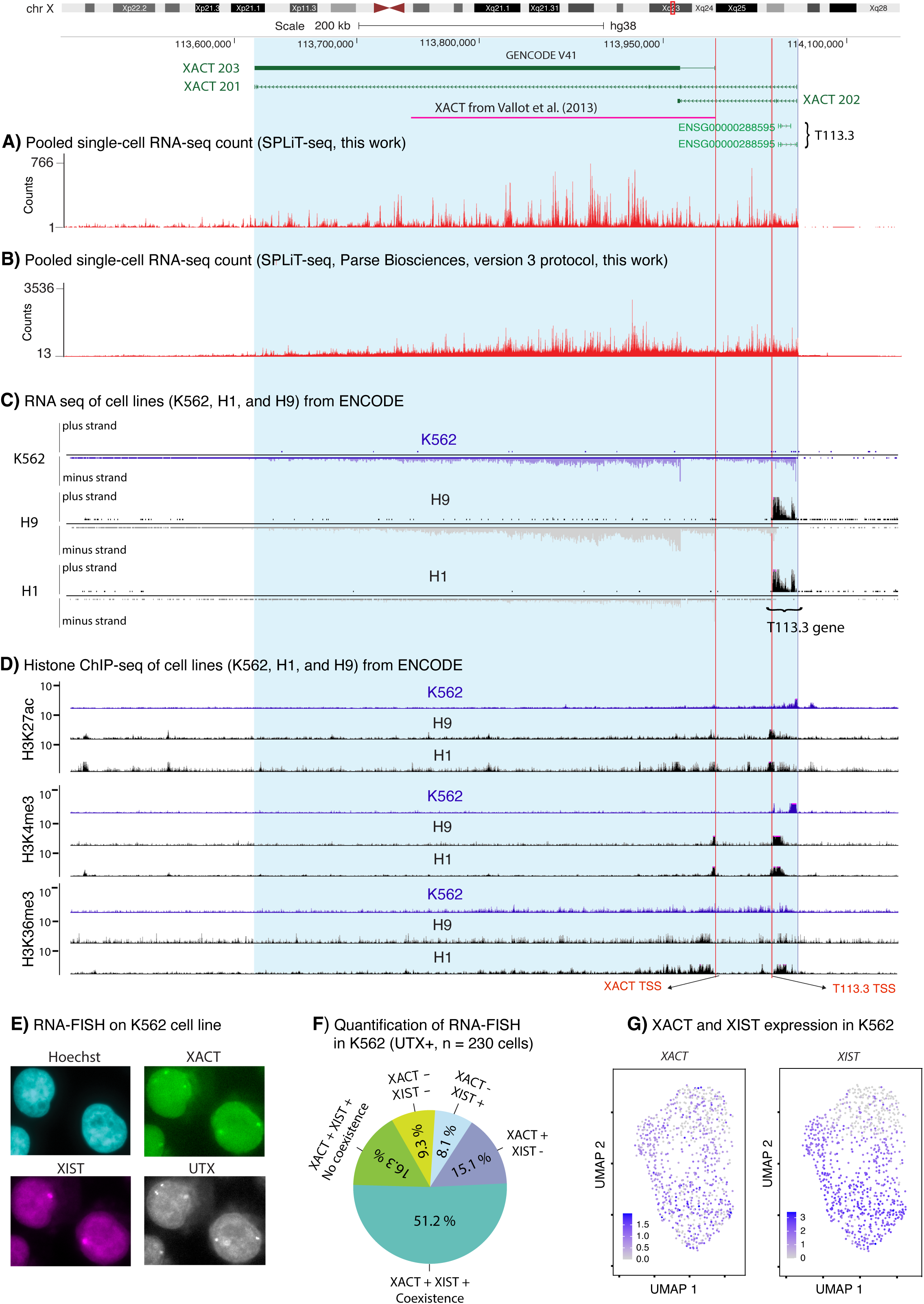
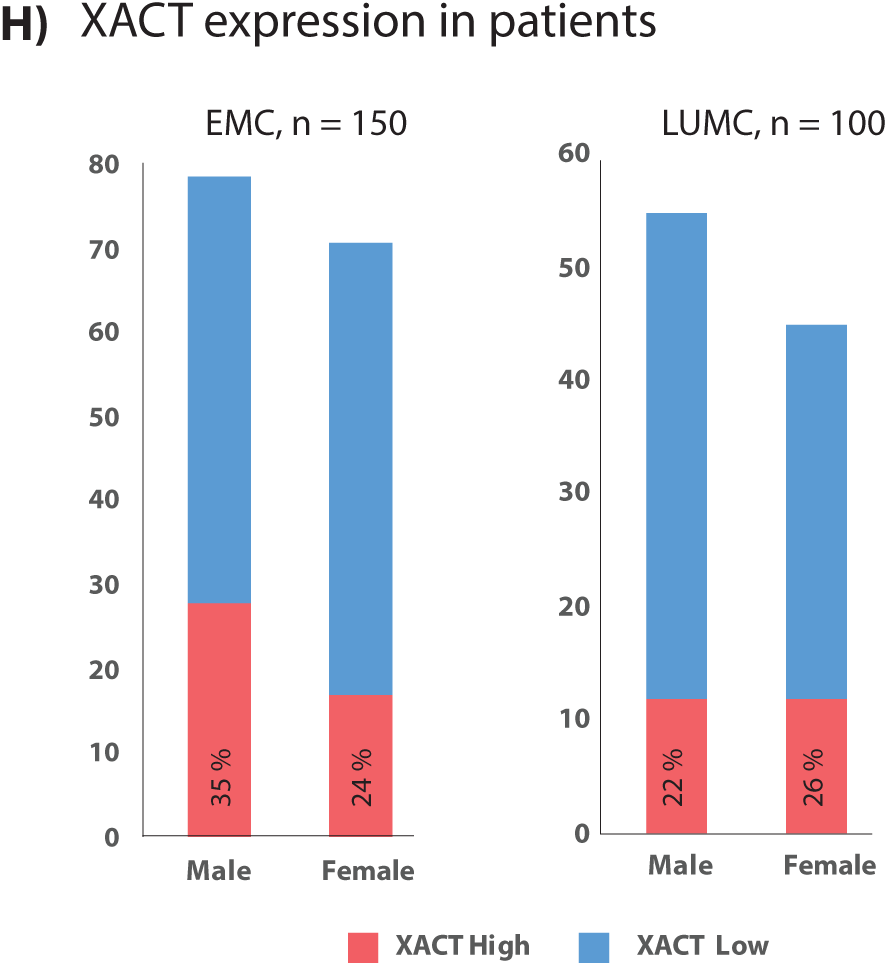
Expression and epigenetic status of the genomic locus encompassing *XACT*. **A)** Visualization of *XACT* lncRNA counts from pooled scRNA-seq in K562 cells; these data were generated using SPLiT-seq in this study. The total RNA-seq counts of all cells are shown on the y-axis. The top panel shows the relevant region of the human genome (GRCh38; chrX:113,458,727-114,143,714). The span of the newly identified *XACT* transcript is depicted in blue throughout panels A-D (GRCh37_chrX:112,859,587-113,302,310, GRCh38_chrX:113,616,300-114,059,121). Several *XACT* transcripts reported in the Ensembl database are shown in green and the 252 kb-long *XACT* transcript originally described in Vallot et al., 2013 ^22^ is indicated in pink (GRCh37_chrX:112,983,323-113,235,148; GRCh38_chrX:113,740,044-113,991,913). The latter is located upstream of another X-encoded lncRNA (*T113.3*), which is expressed from the plus strand ^22^. The *T113.3* gene is located downstream of *XACT* and overlaps with sequences encoding our novel *XACT* transcript. The suggested transcriptional start sites (TSS, also marked throughout panels A-D) of *XACT* and *T113.3* are marked with vertical red lines (*XACT* TSS and T113.3 TSS, respectively). **B)** *XACT* transcript levels in K562 cells, generated using the Parse Biosciences kit and protocol. **C)** Strand-specific total RNA-seq data visualization of the K562 (top graph) and hESC lines (H9, female, and H1, male; middle and bottom graphs). The data were obtained from ENCODE and visualized using the UCSC browser. Signals of unique reads that map to the minus (downwards, below the line) and plus strand (upwards, above the line), respectively, are displayed. **D)** The epigenetic status of the *XACT* locus in K562 and hESC (H1, H9) cells. Histone-3 (H3) ChIP-seq data (for H3K27ac, H3K4me3, H3K36me3, respectively) was obtained from ENCODE, and displayed using the UCSC genome browser. Pooled fold-change of ChIP-seq experiments (from replicates) over control is depicted on the Y-axis. **E)** RNA-fluorescence *in situ* hybridization (RNA-FISH) on K562 cells used to perform SPLIT-seq in panel A. Representative images are shown here, and quantification of all cells is displayed in panel F. **F)** Quantification of RNA-FISH experiment shown in panel E. **G)** Visualization of *XACT* and *XIST* expression in single cells (SPLiT-seq in K562 cells, see panel B). **H)** Expression of *XACT* in AML patients. High (in red) versus low (blue) expression of *XACT* in the EMC and LUMC AML data sets for a total of 250 patients, each split in male and female patients.

Because *XACT* was thought to be expressed during early development ^22,23^, we initially suspected its expression in K562 cells was an artifact. We therefore confirmed our observation through multiple independent experiments: performing another scRNA-seq using an improved SPLiT-seq protocol ^30^ (version 3, Parse Biosciences, Figure 1B); analyzing publicly available bulk RNA-seq data generated from K562 cells (Dataset ERR688 ^31^, Supplementary Figure 1B); comparing *XACT* expression between K562 cells and hESC lines (male, H1 cells; and female, H9 cells; in which *XACT* lncRNA was originally discovered), using ENCODE bulk RNA-seq datasets ^32^ (Figure 1C). In hESCs *XACT* is expressed from a much larger genomic region than originally reported ^22^ (Figure 1C). Notably, in hESCs, *XACT* is expressed from the minus strand, while the nearby lncRNA T113.3 (downstream of *XACT*) is transcribed from the plus strand (Figure 1C). However, in K562 cells *XACT* extends to the *T113.3* gene in the minus strand (Figure 1C), with no expression of *T113.3* on the plus strand. Additionally, we investigated the epigenetic signature of the *XACT* locus, using ENCODE datasets ^32^ (H3K27ac, which marks active promoters and enhancers; H3K36me3, which correlates with transcribed regions; and H3K4me3, which is deposited on active promoters) ^33^. High H3K27ac and H3K4me3 signals were observed near the start site of the 443 kb-long transcript in K562 cells (Figure 1D; Supplementary Figure 2), and H3K36me3 marks covered the gene body. In H1 and H9 cells *XACT* originated from a different start position (Figure 1D; Supplementary Figure 2), suggesting differential promoter usage between K562 cells and hESCs.

Furthermore, we visualized *XACT* expression in K562 cells using RNA-fluorescence *in situ* hybridization (RNA-FISH), with probes for *XACT* ^22,23,27^, and probes detecting *XIST* and UTX (XCI escapee gene) transcripts. This also confirmed *XACT* expression in K562 cells (Figure 1E). Our K562 cells are tetraploid, hence we detected more than two signals for *XACT*. The expression of *XACT* originates from both the Xa and Xi, unlike *XIST* (which is restricted to Xi), and shows significant coexistence with *XIST* on the Xi (Figure 1E,F). Most of the cells (67.4%) expressed all three of these genes, and a few (9.3%) exclusively expressed *UTX*, both *UTX* and *XACT* (15.1%) or *UTX* and *XIST* (8.1%). In cells that expressed these three genes, *XACT* is expressed with *XIST* on the Xi in 75.86%. This was supported by scRNA-seq data from K562 cells where *XIST* is expressed in nearly all cells where *XACT* is expressed (Figure 1G).

Next, since the K562 cell line was derived from a patient with erythroleukemia ^34^, we investigated *XACT* expression in AML patients using two independent RNA-seq datasets of 250 individuals (100 in the LUMC ^35^, and 150 in the EMC ^36^ cohorts). In the EMC cohort, high *XACT* expression was found in 45 patients (30% of the total: 28 males and 17 females, Figure 1H; for the definition of high vs. low expression, see Supplementary Figure 3A). In the LUMC cohort, we found 24 patients with *XACT* expression (24% of the total: 12 males and 12 females, Figure 1H, Supplementary Figure 3A). The length of the *XACT* transcript and its starting point is similar to that seen in K562 cells (Supplementary Figure 3B). However, in some individuals, an even longer transcript was detected (Supplementary Figure 3B). *XACT* expression was validated by RT-qPCR on selected patient samples, and the expected levels of *XACT* were observed (Supplementary Figure 3C).

Thus, we have discovered a novel *XACT* transcript that is expressed in differentiated cells and confirmed its expression in various orthogonal ways. Its expression is further supported by epigenetic profiles of the locus, and distinctively differs from the *XACT* transcript that is expressed in hESCs. The new *XACT* transcript is expressed from both the Xa and Xi and coexists with XIST on the Xi. In addition, this novel *XACT* transcript is also expressed in a subset of male and female AML patients.

### *XACT* expression in AML patients does not lead to decreased expression of *XIST* and reactivation of the inactive X-chromosome

During early-embryonic XCI or during erosion of XCI, *XACT* was proposed to antagonize *XIST’*s silencing activity ^22,23,27^. To understand the consequences of *XACT* reactivation, we used the AML datasets and investigated the impact of *XACT* expression on *XIST*, dosage compensation, and genes that are silenced by XCI, respectively.

First, to investigate the impact of *XACT* expression on *XIST*, we plotted their expression levels in 250 AML patients (from the aforementioned LUMC and EMC datasets) and found no correlation between *XACT* reactivation and *XIST* expression in both male and female AML patients (Figure 2A; Supplementary Figure 4A). Second, we tested whether *XACT* reactivation affects XCI and dosage compensation, despite not having direct influence on *XIST*. The mammalian dosage compensation mechanism maintains an X-chromosome to autosome expression ratio (X:A ratio) of 1 ^37-40^. When we calculated the X:A ratio in AML patient datasets (for LUMC and EMC) with high and low *XACT* expression, we did not observe significant changes. The X:A ratio varied between 0.75 and 1, depending on the origin of the dataset (Figure 2B; Supplementary Figure 3B), indicating that *XACT* expression does not lead to a significant global change in X-linked gene expression levels. Further, we plotted the average expression of X-linked genes (separately for males and females) in relation to high and low levels of *XACT* expression (Figure 2C,D; Supplementary Figure 4C,D) and took chr3 and chr8 (i.e., of similar length and gene content as the X-chr) as controls (Supplementary Figure 5). We observed a slight increase in of X-linked genes expression in *XACT*-high patients, but a similar trend was seen on other chromosomes, indicating that *XACT* reactivation does not lead to specific global upregulation of X-linked genes in both males and females.

**Figure 2.**
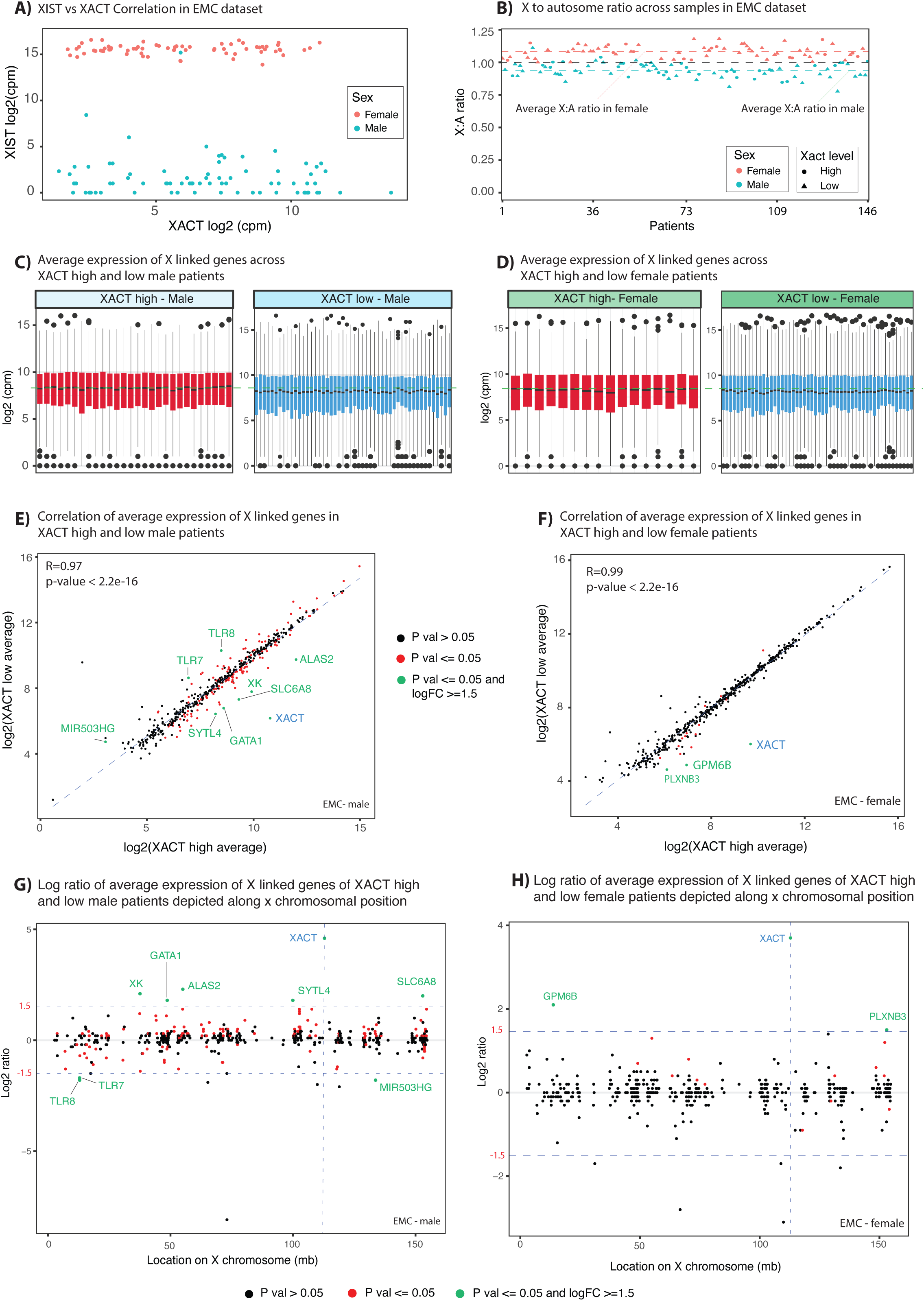
*XACT* expression correlation with *XIST* expression, and impact on dosage compensation. **A)** *XACT* and *XIST* expression correlation in the AML EMC dataset. The *XACT* and *XIST* expression levels (as log2 counts per million, cpm) are depicted on the X- and Y-axis, respectively. Male patients are plotted in turquoise, females in reddish-orange. **B)** X-chromosome to autosome ratio (X:A) of the EMC AML cohort. The *XACT* expression level in each patient is defined as high (dot) or low (triangle) for males (turquoise) and females (reddish-orange), and the average for each sex is indicated by a correspondingly colored dashed line. **C, D)** Boxplots showing average expression of X-linked genes in *XACT* high (red) and *XACT*-low (blue) AML patients, separating males (C) from females (D). **E, F)** Correlation of the average expression of X-linked genes in *XACT*-high and *XACT*-low AML patients, for males (E) and females (F). Genes that show a statistically significant change between *XACT*-high and *XACT*-low (p value <=0.05, T-test) are colored and indicated by red and green dots. Green colored dots and gene names show genes with significant change and a log2 fold change |log2FC| >=1.5. The average expression level in *XACT*-high and *XACT*-low patients is shown on the X- and Y-axis, respectively. **G, H)** Changes in average expression of X-linked genes across the X-chr. Average expression of each gene in *XACT*-high and *XACT*-low AML patients is used to calculate the log2 ratio of each gene that is depicted on the Y-axis. The location of the gene on the X-chr is shown on the X-axis in megabases (mb). The location of *XACT* is shown with a blue vertical dashed line. Males are shown in G, females in H. Genes are colored following the same criteria used in E,F.

We complemented this observation by calculating the average expression of each X-linked gene in *XACT*-high and *XACT*-low patients, which revealed a very high correlation between the expression of X-linked genes between the two groups, again independently of their sex (Figure 2E,F; Supplementary Figure 4E,F). This again indicated that the reactivation of *XACT* does not lead to global changes in the expression of X-linked genes. However, we did observe several genes with significant deregulation (p-value <=0.05, and log2-fold change |log2FC| >=1.5) between *XACT*-high and low AML patients. These include male-specific downregulation of *TLR8* and *TLR7*, which are important for neutrophil activation ^41,42^, as well as male-specific upregulation of genes such as *GATA1*, *ALAS2* and *XK. GATA1* encodes a critical TF in normal and cancerous hematopoiesis, crucial for the healthy differentiation and maturation of erythroid and megakaryocytic lineages ^43,44^. *ALAS2* encodes the first and rate-limiting enzyme in the heme biosynthetic pathway in erythroid cells ^45^. Next, we examined whether these deregulated genes cluster to a specific region of the X-chromosome by plotting them with their annotated transcription start site (TSS) along the X-chromosome (Figure 2G,H; see also Supplementary Figure 3G,H). This analysis did not show clustering of X-linked genes with significant differential expression between *XACT*-high and *XACT*-low AML patients.

We conclude that *XACT* expression does not lead to downregulation of *XIST*, reactivation of genes otherwise silenced by XCI, global upregulation of X-linked genes, or change the X:A expression ratio. However, the reactivation of *XACT* in AML affects the expression of few X-linked genes that have important function in normal and malignant hematopoiesis. These findings suggest that in AML, and perhaps other cancers, *XACT* may have different functions not related to its proposed function in XCI.

### *XACT* is exclusively expressed in erythroid-megakaryocytic lineages during HSC differentiation

We next examined if and when *XACT* is expressed during normal hematopoiesis, the process whereby HSCs differentiate into multipotent progenitors (MPPs), which in turn further develop into intermediate progenitors, including common myeloid progenitors (CMPs) and lymphoid-primed multipotent progenitors (LMPPs). LMPPs generate lymphocytes through common lymphoid progenitors (CLPs), while CMPs give rise to megakaryocytes and erythrocytes via MEPs, and granulocytes and monocytes via GMPs (for reviews, see ^46,47^).

To investigate the expression of *XACT* in this differentiation, we first reanalyzed bulk expression and chromatin accessibility data that were each generated from healthy donors (i.e. from bone marrow and peripheral blood) for 13 human primary blood cell types, thereby spanning the hematopoietic hierarchy (Figure 3A) ^48^. Surprisingly, this analysis showed that *XACT* expression gradually increases, as HSCs differentiate into MPPs and CMPs. It peaks in MEPs and then drops when cells progress towards erythrocytes (Figure 3A,B; Supplementary Figure 6A). Remarkably, *XACT* expression is restricted to the MEP branch and absent in other progenitors (i.e. LMPPs and CLPs, GMPs) and their derivatives. *XACT* shows similar dynamic expression changes in both male and female donors (Figure 3B; Supplementary Figure 6A), while *XIST* expression remained high in all blood cell types of female donors and low in male donors (Figure 3C). The chromatin accessibility data showed that the promoter region of *XACT* becomes accessible only in the MEP branch (starting with HSC, MPP, CMP) of hematopoietic differentiation, which is not observed in all other progenitors and their derivatives, except for B-cells (Supplementary Figure 6B).

**Figure 3.**
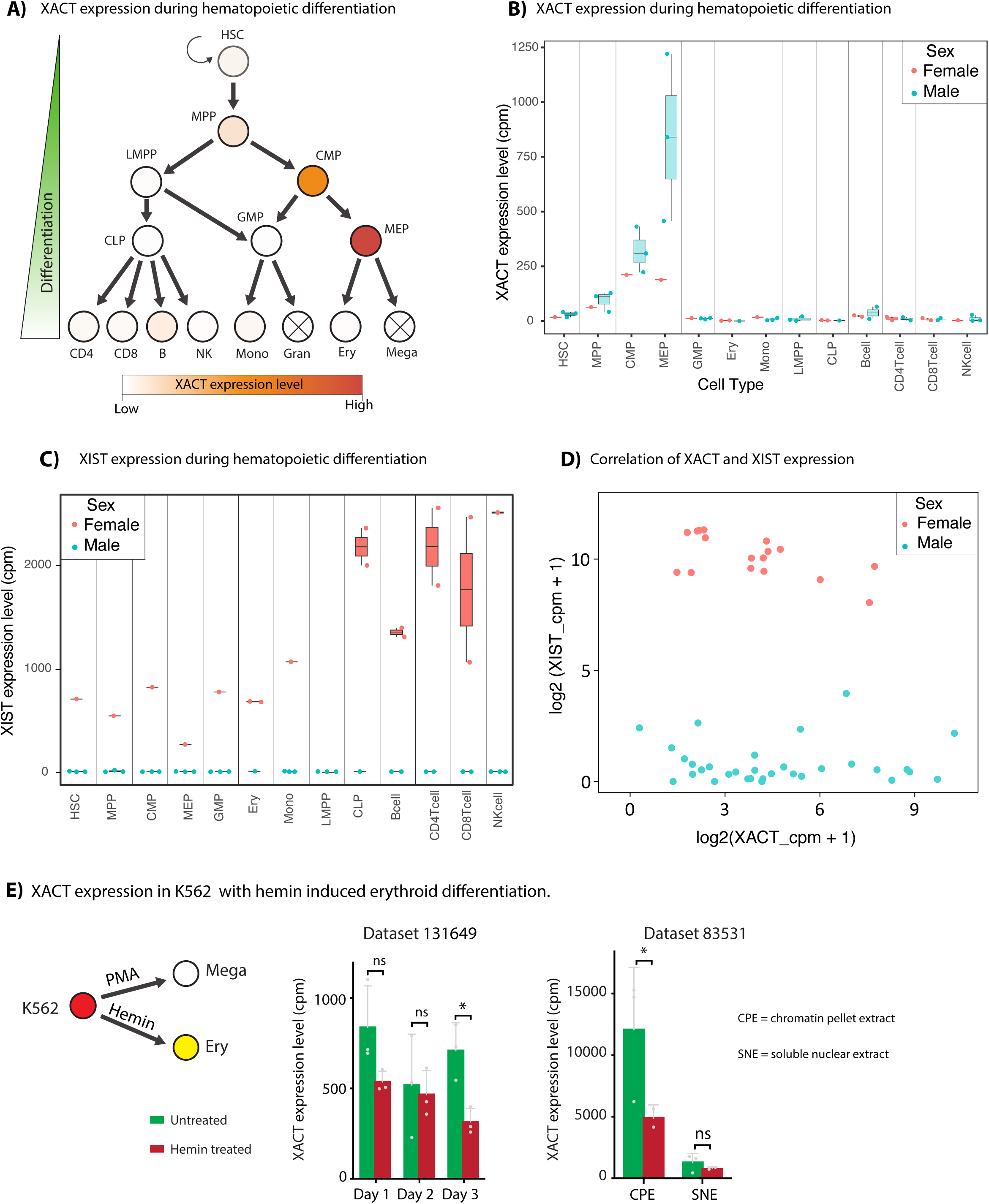
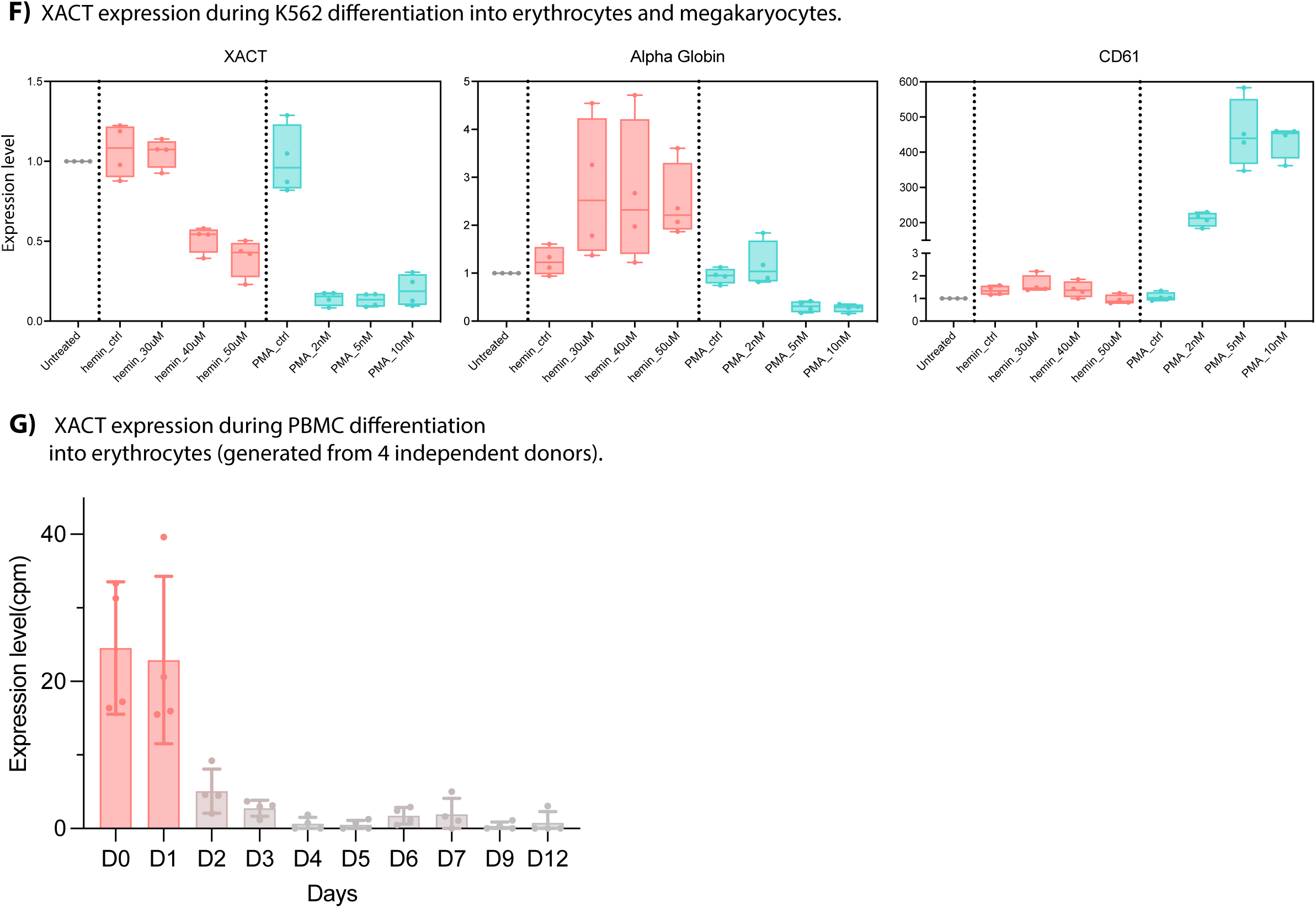
*XACT* expression dynamics during normal hematopoietic cell differentiation. **A)** A simplified hematopoietic cell differentiation diagram (adapted from ^48^) and expression of *XACT*. The *XACT* RNA level is given in color code (brick red, very high expression; orange, low; white, very low; white with cross, not quantified). **B)** Boxplots of *XACT* expression level during normal hematopoiesis. For each cell state/subpopulation (in x-axis), the expression of *XACT* is plotted as counts per million (cpm, y-axis). *XACT* levels are given for males (in turquoise) and females (orange). **C)** Boxplots of *XIST* level during hematopoiesis. For each cell state/subpopulation (x-axis), the expression of *XIST* is plotted as counts per million (cpm, y-axis). **D)** The correlation of *XACT* and *XIST* RNA levels in male (in turquoise) and female (orange) donors. Expression of *XACT* and *XIST* in log2 scale is depicted on the x- and y-axis, respectively. **E)** *XACT* expression in K562 cells subjected to hemin-induced erythroid differentiation (see Materials and Methods ^50,51^). **F)** The expression of *XACT*, alpha globin (used as erythroid lineage marker gene), and the gene *ITGB3* (encoding CD61, a megakaryocytic lineage marker) was quantified in independent samples using qPCR. qPCR was performed after the differentiation of K562 to erythroid and megakaryocytic lineages (using hemin and PMA for 72 hours, respectively). **G)** *XACT* expression during PBMC differentiation into erythrocytes (generated from 4 independent donors, data obtained from ^52^).

Similar to AML patients, in these healthy hematopoietic populations, we observed no correlation between *XACT* reactivation and *XIST* expression of both sexes, (Figure 3D). Additionally, we found that the X:A ratio in the different cell populations is close to 1 (Supplementary Figure 7A). This further supports the notion that *XACT* expression does not affect dosage compensation, and uncouples *XACT* expression from its proposed function in XCI. We further validated the *XACT* expression dynamics by acquiring additional data ^49^ from FACS-sorted HSCs and progenitors (CMPs, GMPs and MEPs) from five healthy donors (4 males, 1 female). This confirmed that *XACT* expression increases during differentiation, peaks in MEPs, and is absent in GMPs (Supplementary Figure 7B).

To assess the effects of MEP differentiation on *XACT* expression, we obtained two RNA-seq datasets generated following hemin-induced erythroid differentiation of K562 cells (GSE131649 ^50^, GSE83531 ^51^). In both we observed a significant downregulation of *XACT* upon cell differentiation (Figure 3E). In the GSE83531 dataset, which was generated after biochemical fractionation of nuclei to isolate chromatin-attached and soluble RNA populations ^51^, *XACT* expression and its decrease upon differentiation are only observed in the chromatin-attached fraction (Figure 3E). To validate these observations, we differentiated K562 cells towards erythrocytes (using hemin) and megakaryocytes (using phorbol myristate acetate, PMA), and performed RT-qPCR for the *XACT* transcript as well as for erythrocyte and megakaryocyte marker genes. We found a significant decrease in *XACT* and increase in α-globin (erythrocyte marker) and CD61 (megakaryocyte marker) (Figure 3F). We further validated this observation by using RNA-seq data generated from pure erythroblast cultures derived from peripheral blood mononuclear cells (PBMC) donors ^52^ (these PBMCs were cultured and expanded for 10 days, followed by 12-day differentiation into erythrocytes ^52^). In this setting, *XACT* was highly expressed in the erythroid progenitors and quickly downregulated between Day 1-2 of differentiation (Figure 3G).

Thus, *XACT* is expressed specifically in the megakaryocyte-erythrocyte lineage specification in both males and females. *XACT* expression is linked to epigenetic changes of the *XACT* locus and is not correlated with *XIST* expression. Upon differentiation of MEPs, the expression of *XACT* is significantly downregulated, the promoter becomes closed accordingly, and *XACT* RNA is found chromatin-bound. These results suggest that the XACT lncRNA serves other functions beyond its role in XCI.

### *XACT* is important to maintain MEP proliferation as well as differentiation

To understand the function of *XACT* in hematopoiesis, we inactivated *XACT* expression (using CRISPRi) in HUDEP2 cells (an MEP-like cell line with erythroid differentiation potential), where *XACT* is highly expressed and downregulated upon erythroid differentiation (Supplementary Figure 8 A,B). First, we compared the growth rate of HUDEP2-KD (i.e. *XACT*-inactivated) and HUDEP2-NT (*XACT*-active) cells, during proliferation and after induction of differentiation. Proliferation of HUDEP2-KD cells was slower compared to HUDEP2-NT (Figure 4A). Upon differentiation, the HUDEP2-KD cells showed significant cell death and after 3 days most of the cells had died, whereas the HUDEP2-NT cells progressed without survival problem, as reflected by the amount of RNA obtained (Supplementary Figure 8C). Second, we performed RNA-seq on proliferating HUDEP2-KD and HUDEP2-NT cells, as well as after induction of their erythrocyte differentiation (i.e. at day (D) 3). Next, to deconvolute the bulk RNA-seq data with CIBERSORTx ^53^, we first generated specific gene expression profiles corresponding to different differentiation time points of erythroblasts (GSE124363). These profiles were then used as a reference to estimate the composition of HUDEP2 cells under both proliferation and differentiation states using CIBERSORTx. This analysis confirmed the start of premature differentiation, characterized by a decrease in both proliferating and differentiating HUDEP2-KD cells, which remained at D0-4 stages, and an increase in cells progressing beyond these initial stages. Following this, the result showed the blockage of differentiation and/or cell death, reflected in differentiating HUDEP2-KD cells mainly remaining at D5-7 stages, and fewer such cells advancing to further differentiation stages at D9-12 stages (Figure 4B).

**Figure 4.**
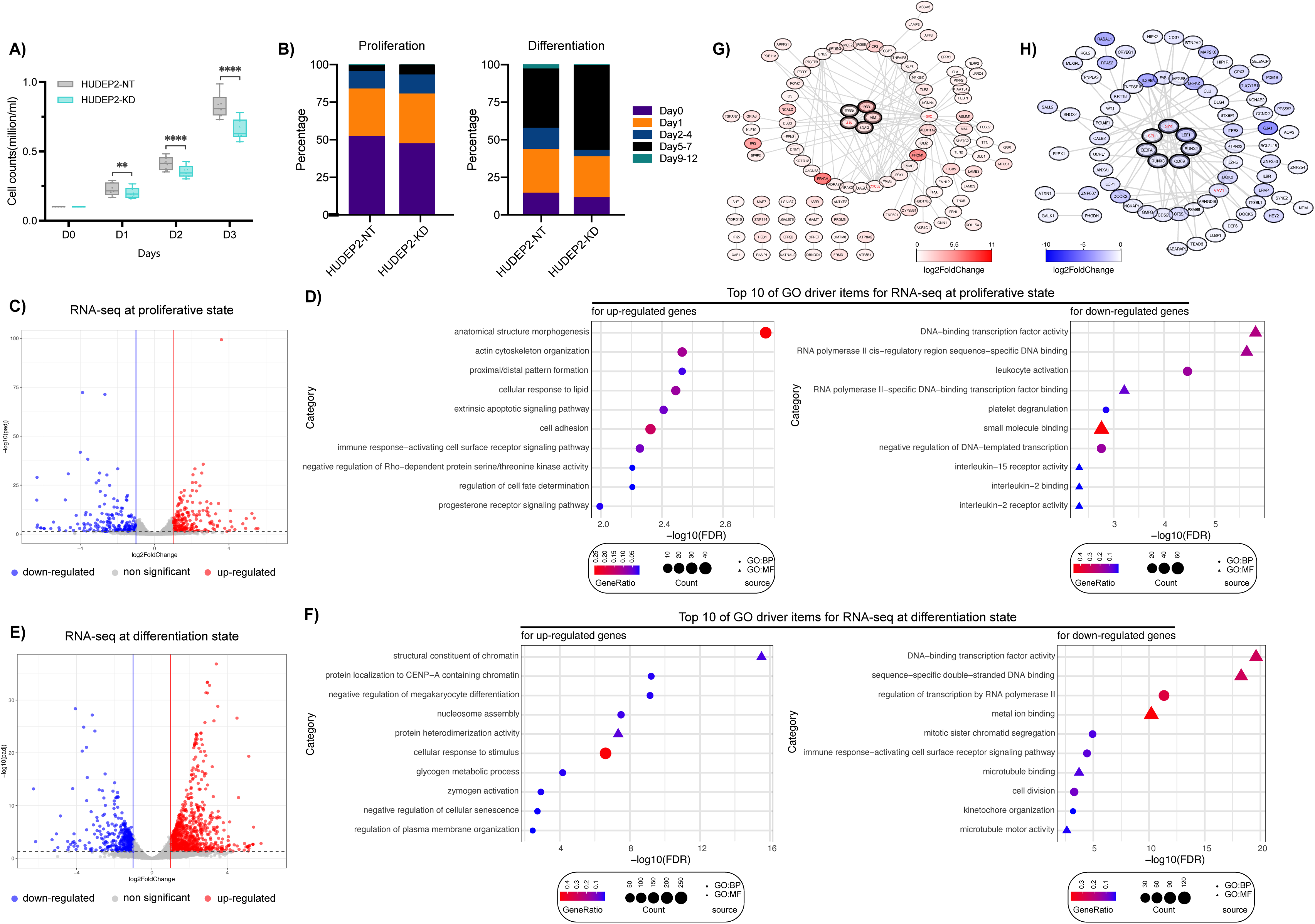
Impact of *XACT* inactivation on proliferation, differentiation, and global gene expression in HUDEP2 cells. **A)** Proliferation of HUDEP2-KD (turquoise) and HUDEP2-NT (gray) cells. **B)** Cell type abundances of HUDEP2-KD and HUDEP2-NT cells estimated from bulk RNA-seq, at proliferating cell stage and after induction of erythroid differentiation (at D3; see Materials and Methods). **C)** Differentially expressed genes (DEGs) between HUDEP2-KD and HUDEP2-NT during cell proliferation. **D)** Biological process (GO:BP) and Molecular Function (GO:MF) GO-terms enriched in up and down regulated genes upon *XACT* inactivation, under proliferation condition (see also panel C). **E)** DEGs between HUDEP2-KD and HUDEP2-NT identified after 3 days of differentiation. **F)** GO:BP and GO:MF GO-terms enriched in up and down regulated genes upon *XACT* inactivation after 3 days of differentiation (Figure 4E). **G)** Protein-protein interaction (PPI) networks derived from highly-expressed genes in proliferating HUDEP2-KD cells. Genes with bold outlines belong to the most densely connected module, while genes marked in red represent the top-3 hub nodes. **H)** PPI networks derived from lowly-expressed genes in proliferating HUDEP2-KD cells. The same outlines and colours were used as in panel G.

We next identified the differentially expressed genes (DEGs; with padj <= 0.05; and |log2FC| >= 1) between HUDEP2-NT and HUDEP2-KD groups. In the proliferative state, we obtained 205 down and 253 upregulated genes (Figure 4C; Supplementary Table 2). These first 205 downregulated genes in HUDEP2-KD were enriched for gene ontology terms (GO terms) related to TF DNA−binding activity and *cis*−regulatory binding (for the complete list in Figure 4D, see Supplementary Table 2). The 253 upregulated genes are associated with actin cytoskeleton organization at cellular level, and organismal anatomical structure morphogenesis and proximal/distal patterning (Figure 4D; Supplementary Table 2). In the differentiation state, RNA-seq data DEG analyses identified 409 down and 942 upregulated genes (Figure 4E; Supplementary Table 3). The 409 downregulated genes in the HUDEP2-KD were enriched for GO terms similar to those identified for proliferating cells (Figure 4F; Supplementary Table 3), indicating a possible role of *XACT* in regulation of transcription. In addition, we also obtained terms related to mitotic sister chromatid segregation, cell division, and kinetochore organization (Figure 4F; Supplementary Table 3), suggesting possible ways leading to cell death upon differentiation of HUDEP2-KD cells.

To identify key components in the *XACT*-dependent gene network, we constructed protein-protein interaction (PPI) networks involving the up and downregulated genes, comparing HUDEP2-KD with HUDEP2-NT proliferating cells. For upregulated genes in HUDEP2-KD cells, the PPI network analysis (using Molecular Complex Detection, MCODE ^54^) identified a significant module that included the genes *VIM*, *SNAI2*, *JUN*, *PGR*, and *ERBB4*. Among these, the top-3 hub genes (calculated by cytoHubba ^55^ based on degree) were *SRC*, *JUN*, and *CXCL8* (Figure 4G; Supplementary Table 4) of which *JUN* has been reported to act as a positive regulator to induce cell quiescence ^56,57^. For downregulated genes in HUDEP2-KD cells, the significant module included *SPI1* (*PU.1*), *SYK*, *LEF1*, *CEBPA*, *RUNX2*, *RUNX3*, and CD69, with the top hub genes being *SYK*, *SPI1*, and *VAV1* (Figure 4H; Supplementary Table 4). Notably, SPI1-deficient erythroid precursors exhibit loss of self-renewal, impaired proliferation, premature differentiation, and cell death ^58^.

We conclude that *XACT* plays a crucial role in controlling the balance between proliferation and differentiation of erythrocyte progenitors, and that its absence affects several TFs involved in these processes, leading to delayed proliferation, failure to mature into erythrocytes, and cell death.

### *XACT* is important for recruiting regulatory element binding proteins and for transcriptional regulation

To better understand the functions of *XACT*, we employed a multi-omics approach, assessing chromatin accessibility (ATAC-seq), H3K27ac and H3K27me3 profiles (Cut&Tag), and RNA-seq, comparing HUDEP2-KD and HUDEP2-NT cells (see Supplementary Note 1 for details).

In the ATAC-seq analysis, we identified 685 differentially accessible regions (DARs) (|log2FC| >= 1 and FDR <= 0.05) with higher accessibility and 741 regions with lower accessibility in HUDEP2-KD relative to HUDEP2-NT (Figure 5A; Supplementary Figure 8D; Supplementary Table 5). These DARs are located at different sites across the genome, such as at promoters, introns, and distal intergenic regions, and they showed no preference for the X-chromosome (Figure 5B, Supplementary Figure 8G). DARs with high enrichment in HUDEP2-KD cells are fewer at promoter regions (within 1 kb, Figure 5B). TF-occupancy (footprint) prediction (using TOBIAS ^59^) identified 39 TFs with a higher occupancy in HUDEP2-KD cells, and 43 TFs with higher occupancy in the HUDEP2-NT cells (Figure 5C; Supplementary Table 5). The TFs with lower occupancy in HUDEP2-KD included ETS family members (e.g., SPI1, SPIB, FLI1, ETV1, ETV4) and other TFs such as TAL1, CEBPA, CEBPB, and CEBPD (Figure 5C; Supplementary Table 5). *SPI1* and *CEBPA are* important for proliferation and differentiation of erythroid progenitors ^58,60-63^. They are downregulated in HUDEP2-KD cells (RNA-seq) and SPI1 was also a hub in our protein network analysis (Figure 4H). The TFs with higher occupancy in HUDEP2-KD cells include the activator protein-1 (AP-1) family of proteins (e.g., JUN, JUNB, JUND, ATF3, BATF, FOSB, FOSL2, and FOS) (Figure 5C; Supplementary Table 5). This is consistent with the protein network analysis wherein we found JUN as a hub and among the up-regulated genes in HUDEP2-KD cells (Figure 4G). AP-1 family proteins play an important role in erythroid differentiation by activating erythroid-specific genes ^64^ ^56,57,65-67^. We also complemented this TF footprint analysis with motif analysis of DARs and, accordingly, found an enrichment for ETS family members (such as SPI1, SPIB, FLI1) in regions with lower accessibility in HUDEP2-KD, and AP-1 family motifs in regions with higher accessibility (Supplementary Table 5).

**Figure 5.**
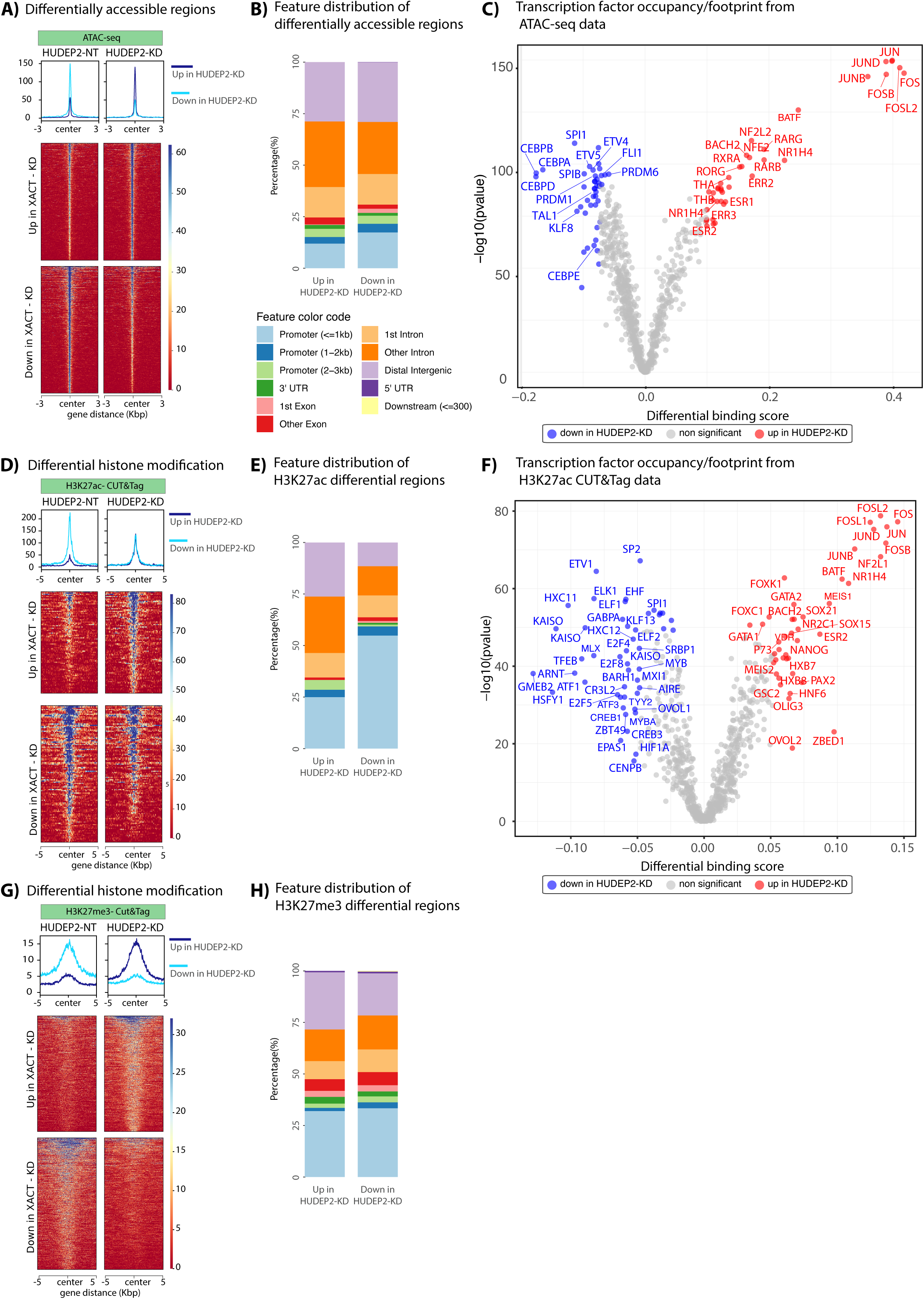
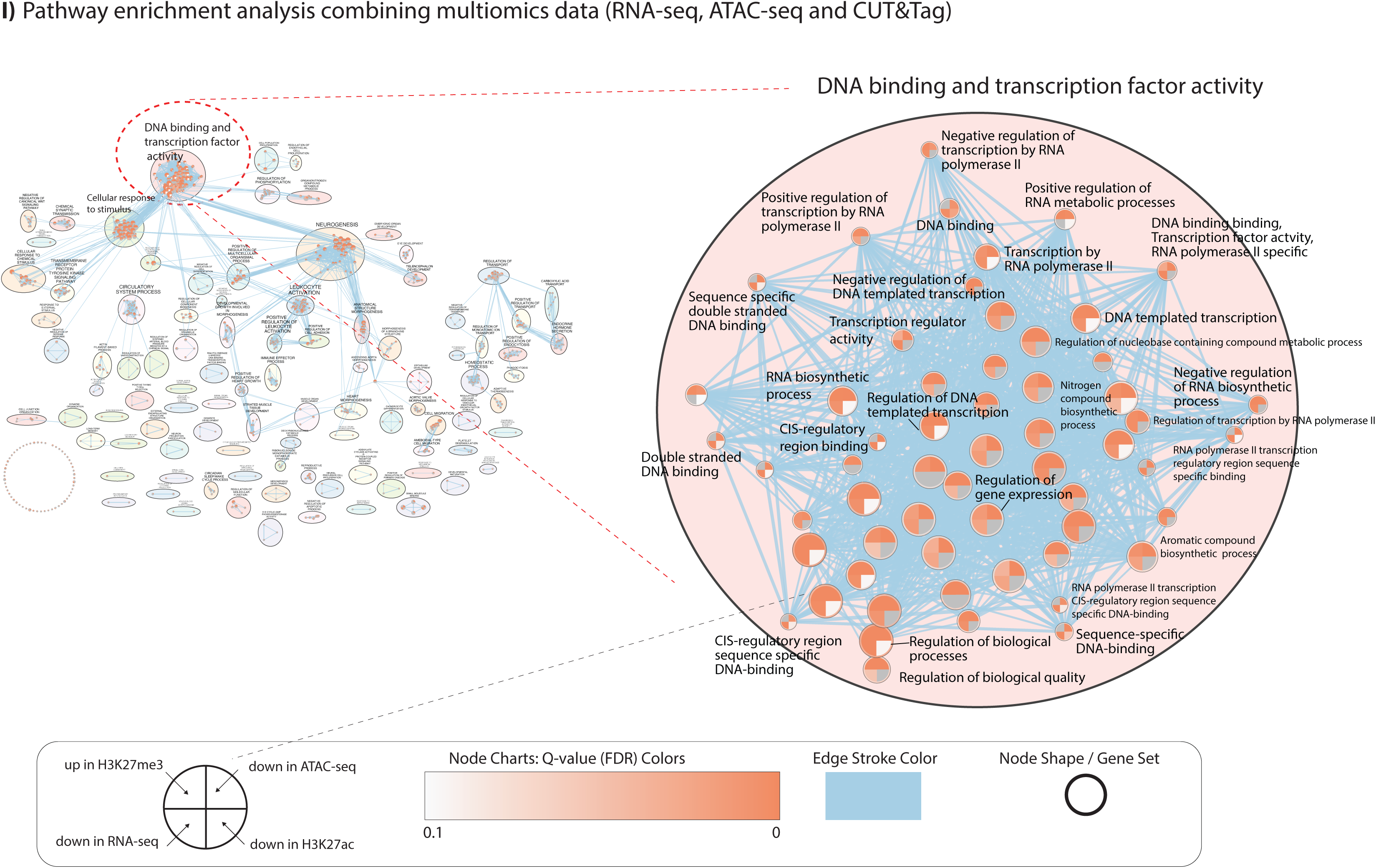
Assessment of *XACT* Functions via Chromatin Accessibility and Histone Signatures in HUDEP2 Cells. **A)** Differentially accessible regions (DARs), up (higher accessibility) and down (lower accessibility), as determined by ATAC-seq, of HUDEP2-NT (*XACT*-positive) and HUDEP2-KD cells (wherein *XACT* is inactivated using CRISPRi). **B)** Feature distribution of DARs over loci and subregions thereof (see colour codes for these features). **C)** Transcription factor (TF) occupancy/footprint derived from the ATAC-seq data. Blue circles indicate TF footprints that show decreased accessibility in HUDEP2-KD cells compared to HUDEP2-NT; red circles indicate increased accessibility. **D)** Differential H3K27ac marks between HUDEP2-NT and HUDEP2-KD cells (obtained using CUT&Tag). **E)** Feature distribution of regions that show differential H3K27ac mark levels (same colour codes as in B). **F)** TF occupancy/footprint analysis results from H3K27ac data. Blue dots indicate TF footprints in regions showing decreased H3K27ac in HUDEP2-KD cells (compared to HUDEP2-NT); red circles indicate increased H3K27ac. **G)** Differential H3K27me3 marks for HUDEP2-NT and HUDEP2-KD cells. **H)** Feature distribution of regions that show differential H3K27me3. **I)** Pathway enrichment analysis combining multi-omics data for Group 1 (genes that show down-regulation (RNA-seq), regions that show low chromatin accessibility (ATAC-seq), low H3K27ac, and high H3K27me3 in HUDEP2-KD cells. The different pathways identified in this analysis are depicted on the left. The most significantly enriched cluster (i.e. DNA-binding and TF activity, named after the most significant node in this cluster in RNA-seq) is depicted on the right. Each node carries information about the data types and significance (in colour code). The connections between nodes (edges) are colored in blue (for more details, see Supplementary Figure 9).

In the H3K27ac CUT&Tag data, we identified 197 regions with significant changes (|log2FC| >= 0.5, FDR <= 0.05) between HUDEP2-KD and HUDEP2-NT (84 with high H3K27ac in HUDEP2-KD, and 113 with low H3K27ac) (Figure 5D, E; Supplementary Figure 8E; Supplementary Table 6), and with no specific preference for the X-chromosome (Supplementary Figure 8G). In the HUDEP2-KD group, sites with decreased H3K27ac mark were frequently located in the promoter region (<=1 kb) (Figure 5E), further pointing to functions of *XACT* in recruiting TFs and hence transcriptional regulation. TF footprint and motif search on these H3K27ac-marked differential binding regions identified similar profiles to that of the chromatin accessibility data (Figure 5F; Supplementary Table 6; see Supplementary Note 1 for details) relating to the functions of *XACT* in co-operation with AP-1 and ETS family TFs.

In the H3K27me3 data, we identified 1243 regions with significant H3K27me3 marks (|log2FC| >= 1 and FDR <= 0.05), with 581 regions showing higher H3K27me3 in HUDEP2-KD compared to HUDEP2-NT, and 662 regions with lower H3K27me3 (Figure 5G; Supplementary Figure 8F; Supplementary Table 7). These differential H3K27me3 marks were distributed throughout the genome (Figure 5H), with no specific enrichment on the X-chromosome (Supplementary Figure 8G). Motif analysis of the -/+ 200 bp-long regions, around the TSS of the genes covered by differential H3K27me3 binding regions (Supplementary Table 7), showed a trend similar to these from the H3K27ac and chromatin accessibility data, further linking *XACT* co-operation with AP-1 and ETS family of TFs (see Supplementary Note 1 for details).

To gain a more comprehensive understanding of the function(s) of *XACT*, we performed pathway enrichment analysis using a large list of gene sets obtained from RNA-seq, as well as of the genes annotated using the chromatin accessibility and histone modification (see Supplementary Note 1 for details). This integrative analysis also showed the role of *XACT* via terms related to DNA-binding and TF activity, *cis-*regulatory region binding, transcription by RNAPol-II, and DNA-templated transcription, supporting the role of *XACT* in recruiting TFs and its role in regulation of transcription (Figure 5I; for more details, Supplementary Figures 9 and 10; Supplementary Table 8).

In conclusion, our multi-omics approach shows that the function of *XACT* extends beyond its effects on the X-chromosome and affects several regions throughout the genome. The absence of *XACT* leads to changes in the recruitment of several TFs important for the proliferation and differentiation of erythroid progenitors. Our results also highlight the importance of *XACT* in recruiting TFs to *cis*-regulatory regions and hence in transcriptional regulation.

### *XACT*’s exclusive expression in the erythroid-megakaryocytic lineage during hematopoiesis can indicate the blast origin in AML patients

The dynamics of *XACT* expression during normal hematopoiesis (Figure 3A,B) suggested a possible role in erythroid-megakaryocytic lineage specification, whereas its absence may promote other lineages potentially reflected in AML patients. To investigate this, we manually curated and selected 98 AML patients for DEG analysis (60 from EMC; 38 from LUMC cohorts) with the highest and lowest *XACT* expression, and validating expression level by visually inspecting read alignment at the *XACT* locus. DEG analysis was performed independently due to differing RNA-enrichment methods (i.e. Ribo depletion vs. polyA enrichment).

In the EMC dataset, we found 641 DEGs (429 DEGs down in *XACT*-high patients, and 212 DEGs up) (Figure 6A). In the LUMC dataset, these numbers were 1196 DEGs with 593 down and 603 upregulated, DEGs; Figure 6A). When we matched the DEGs from these two datasets, 195 down and 88 upregulated genes in common were identified (Figure 6A; Supplementary Table 9). The 195 genes downregulated with high-*XACT* were significantly associated with GO terms related to neutrophil degranulation, neutrophil activation involved in immune response, neutrophil-mediated immunity, and other immune response-related terms (Figure 6A; Supplementary Table 9). In contrast, the 88 upregulated genes were associated with erythroid, myeloid, and megakaryocyte differentiation (Figure 6A; Supplementary Table 9). These findings further support our prior observation of *XACT*’s specific expression and possible function of *XACT* in erythroid lineage differentiation. The genes that are upregulated with *XACT* upregulation included those encoding the TFs TAL1, GATA1, and KLF1, which are crucial for the differentiation into erythroid and megakaryocytic cell lineages ^46^ ^44,68,69^. In the list of genes downregulated in *XACT*-high AML patients, the toll-like receptor (TLR) pathway is enriched, which is important in the innate immune system and in neutrophil activation ^41,70^.

**Figure 6.**
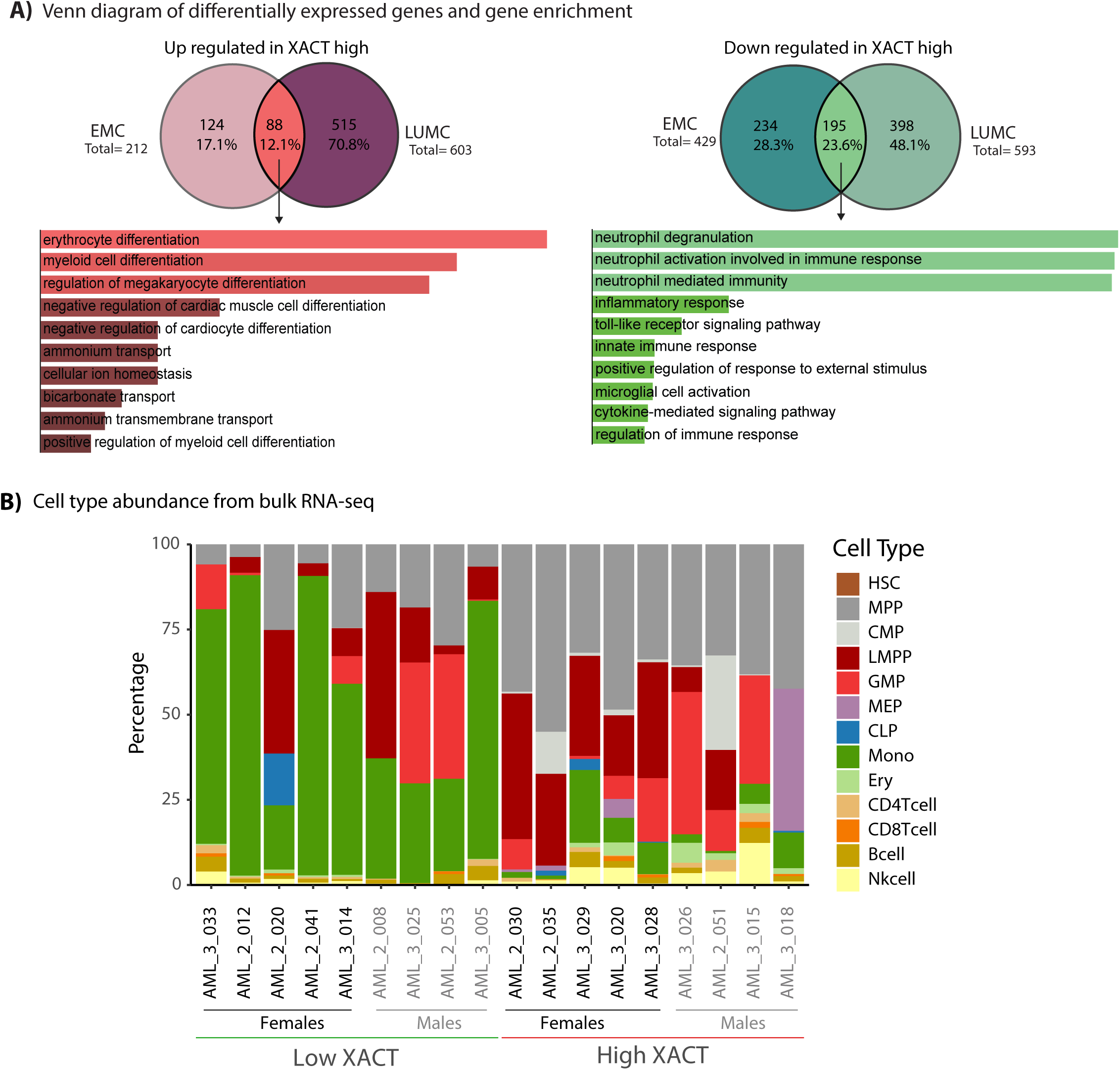
*XACT* expression in AML is associated with a distinct cell of origin. **A)** Venn diagram and gene enrichment analysis (GEA) on differentially expressed genes (DEGs) between patients with high and low *XACT* levels. For the EMC dataset, 60 patients with *XACT*-high and low expression are used (30 high-*XACT*; 30 low). For the LUMC dataset, 38 patients with *XACT*-high and low expression are used (19 high-*XACT*; 19 low). Genes with |log2FC| > 1, and adj.pvalue <0.05 are defined as DEGs. DEGs overlapping between the LUMC and EMC datasets were used for GEA. The number of DEGs in each dataset is depicted next to the Venn diagrams. **B)** Cell type abundance from bulk RNA-seq of *XACT*-high and low AML patients. Normalized counts of bulk RNA-seq data were used as input for CIBERSORTx analysis.

After confirming the lineage-specific expression of *XACT*, we investigated whether AML patients with high and low XACT levels exhibited blasts derived from specific blood cell types to determine whether *XACT* expression could be used as a discriminatory factor for (subclasses of) AML patients. For this, we employed CIBERSORTx ^53^ to estimate the abundance of different cell types from bulk RNA-seq data obtained from AML patients with high and low *XACT*. The results show that AML patients with low *XACT* often had blasts originating from monocytes (Figure 6B), whereas AML patients with high *XACT* frequently exhibited blasts derived from MPPs, MEPs and CMPs (Figure 6B). These findings align with the observed expression dynamics of *XACT* during normal hematopoiesis.

In conclusion, our investigations show that the erythroid-megakaryocytic lineage-specific *XACT* expression and functions during HSC differentiation is also reflected in AML patients. Genes associated with neutrophil-related terms were downregulated in *XACT*-high patients, while genes linked to erythrocyte differentiation were upregulated. Key TFs that play important roles in erythropoiesis (such as TAL1, GATA1, and KLF1) were upregulated with increased *XACT* level. Additionally, *XACT* expression is correlated with the blast differentiation stage observed in AML patients, which was found consistent with *XACT* expression normal hematopoiesis. These findings highlight the potential of *XACT* expression as a discriminatory factor in AML and its role in disease progression.

## Discussion

In this study, we present the discovery of a novel transcript of lncRNA *XACT* in both normal adult cells and cancer cells in males and females. We show that this new *XACT* is expressed from both the Xi and Xa in the presence of *XIST* on the Xi. Intriguingly, *XACT* expression neither impacts expression of *XIST* nor the reactivation of X-linked genes silenced during XCI. During hematopoietic differentiation, *XACT* is specifically expressed in MEPs both in males and females. The highest *XACT* levels are observed in MEPs and diminish in mature megakaryocytes and erythrocytes. *XACT* is likely needed for the self-renewal of MEPs, and blocking differentiation of MEPs into erythrocytes by maintaining the balance between ETS and AP-1 TF-family members. We also show that *XACT* (co-)regulates gene transcription by recruiting TFs to *cis-*regulatory genomic regions. Finally, we document *XACT* expression in subsets of (male and female) AML patients, and report that increased *XACT* levels can be associated with the origin of erythroid-megakaryocytic blasts.

The novel *XACT* transcript we identified is ≈443 kb-long, which is longer and different from the originally described ≈252-kb transcript expressed only in hESCs. Our work corrects the latter finding showing that the original *XACT* transcript is ≈123 kb longer (hence a length of ≈375kb). Additionally, we show a switch in transcript usage between pluripotent cells in the embryo (*XACT*-RNA length ≈375kb) and adult healthy as well as diseased somatic cells (≈443kb) (see model, Figure 7). The novel long *XACT* transcript is generated from an alternative TSS and is marked by distinct histone and chromatin signatures, spanning the entire 443-kb locus, making it one of the longest transcribed genes in the human genome. We propose classifying it as a new type of RNA, such as "extra-long" non-coding RNA (xlncRNA), while emphasizing the need for an improved lncRNA classification system based on function(s), rather than the arbitrary grouping of any non-coding RNA longer than 200 nucleotides.

**Figure 7.**
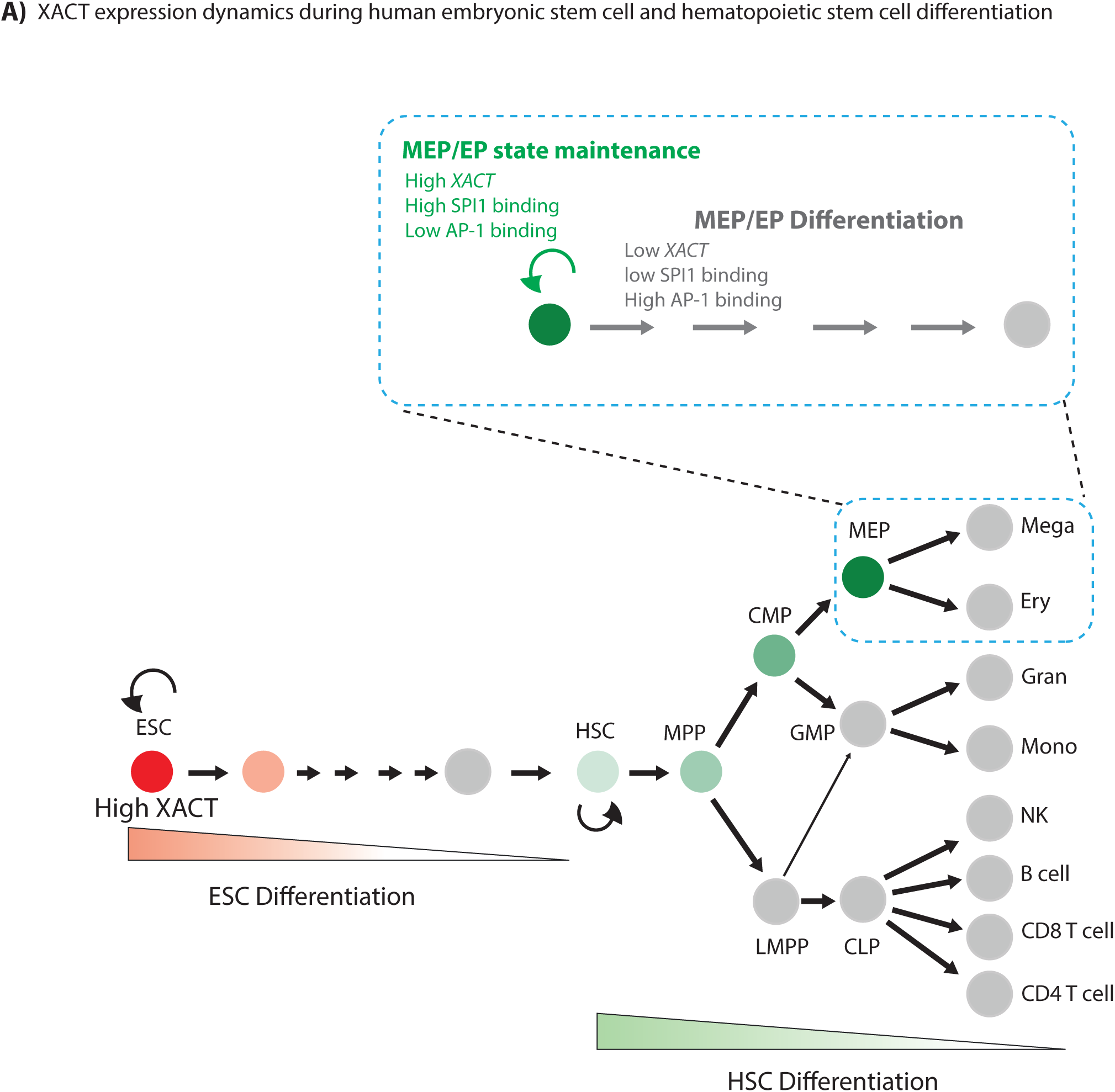
The dynamics of *XACT* expression during early development and hematopoiesis. **A)** Based on the literature and our observations in this study, the dynamics of *XACT* expression in human embryonic stem cells (hESC; see reddish part) and hematopoiesis (greenish) is depicted. Although *XACT* is expressed but quickly down-regulated in hESCs, its functions in these undifferentiated pluripotent cells remain debated (see main text). The novel *XACT* transcript (in green) identified in this study is proposed to exert a specific function during hematopoiesis in erythroid-megakaryocytic lineage specification. Our research identifies *XACT* as a key co-regulator in MEP/erythroid progenitor (EP) state maintenance (green) and subsequent cell differentiation (blue).

During early development, the short *XACT* isoform function has been linked to XCI ^22,23,27^. In the mammalian early embryo, XCI is triggered by the lncRNA *XIST* ^5,11-13^, and *XACT*, which was shown to be expressed from - and coat - the Xa in both males and females. *XACT* was proposed to counter the ability of *XIST* to induce XCI ^22,23,27^; however, these functions of *XACT* in XCI have also been contested ^28^. In one *XACT* deletion study in primed hPSCs, *XACT* was found dispensable for X-linked gene repression on the Xi while influencing neural differentiation^28^. Recently, another *XACT* deletion study in naive hPSCs concluded that *XACT* neither antagonizes *XIST* nor controls X-chromosome activity ^29^, contrary to previous claims ^23^. Our work reveals novel functions of *XACT* in healthy adult somatic and in malignant cells, where its expression neither affects *XIST* nor activates X-linked genes inactivated by XCI, extending the role of the *XACT* beyond the debated function in early embryogenesis.

*XACT* can be added to the list of emerging important lncRNAs in tissue stem cells, including skin, brain, and muscle as well as in cell pluripotency and developmental patterning ^71^. We link *XACT*’s function, to our knowledge for the first time, to hematopoietic lineage specification, providing a solid basis for pinpointing its precise function(s) and mode(s) of action in future research. The exclusive expression of *XACT* in MEPs during HSC differentiation, supported by its temporal expression dynamics (with decreased levels upon differentiation into erythrocytes and mega-karyocytes), strongly points to a potential role in cell fate decisions. While this function of *XACT* is novel, previous work on *Xist*, mainly in mice, has shown the role of *Xist* in hematopoietic cell fate decisions. Deletion of *Xist* in mouse HSCs leads to progressive hematopoietic defects, resulting in hematologic malignancies and lethality in female mice ^72^. *Xist* deletion facilitates lymphoid progenitor differentiation, but hinders myeloid progenitor differentiation towards the megakaryocytes and erythrocytes ^73^. In our analysis of human hematopoietic differentiation, *XIST* expression remains unchanged, and *XACT* exhibits lineage-specific expression in both sexes, leading us to hypothesize that in humans *XACT* may have taken over the some of the functions of *Xist* seen in mice.

In our attempt to understand *XACT*’s mechanism(s) of action, we show that inactivating *XACT* in HUDEP2 cells disrupts proliferation and differentiation, and downregulates genes involved in transcription regulation and *cis*-regulatory region binding, consistent with the general gene-regulatory functions of lncRNAs ^1,2,74,75^. Additionally, the changes in chromatin accessibility and H3K27ac observed upon *XACT* inactivation primarily mapped around the promoter regions of genes, indicating that *XACT* lncRNA acts in *trans* to recruit regulatory proteins. Although mechanisms of lncRNAs are generally poorly understood, current evidence suggests lncRNAs regulate local (in *cis*) and distal (in *trans*) expression of genes by regulating enhancer-promoter interactions, chromatin architecture through direct lncRNA-DNA binding or via protein-mediated DNA-binding ^1,2,74-77^. lncRNAs can bind to histone-modifying complexes, DNA-binding proteins (including TFs), and RNAPol-II ^1,2,74-77^. The specific action of *XACT*, whether through direct DNA-binding or via mechanisms mediated by other proteins or factors, still needs to be established. However, motif and footprint analyses of ATAC-seq and CUT&Tag data, along with protein-protein interaction network analysis (in particular comparing HUDEP2 cells with and without *XACT* expression), revealed a loss of ETS family (e.g., SPI1, FLI1, ETV5) and a gain of AP-1 family TFs (e.g., JUN, JUNB, BATF, FOS) upon *XACT* inactivation, suggesting potential co-operative partners (direct or indirect) of *XACT*.

The ETS family of TFs play diverse roles in cell proliferation and differentiation, cell cycle control, cell migration, and cell death ^78^. For example, SPI1 is essential for myeloid lineage commitment, suppresses erythroid differentiation, and maintains the balance between different hematopoietic lineages and cell fate decisions ^58,60-63^. FLI-1 is vital for megakaryocyte development ^79,80^. The AP-1 family TFs also play important functions in proliferation, differentiation, and apoptosis^66^. In MEP cells, AP-1-type TFs support erythroid differentiation by activating erythroid-specific genes and they promote erythropoiesis by counteracting the repressive effects of factors like PU.1 ^64^ ^56,57,65-67^. ETS and AP-1 family TFs compete for binding to regulatory regions of target genes ^56,57^, potentially influencing cell commitment to either the myeloid or erythroid lineage. In the context of our work, the *XACT* may facilitate SPI1 binding to maintain the MEP (or erythroid progenitor, EP) state. When *XACT* is absent and SPI1 binding is lost, AP-1 factors take over and promote differentiation and cell cycle arrest (see model, Figure 7). However, proper differentiation still requires SPI1 for cell survival, leading to cell death upon differentiation in absence of SPI1.

In conclusion, our study has provided valuable insights into the unique and intricate expression patterns of *XACT* in both healthy and cancer somatic cells, opening new dimensions beyond its proposed exclusive role in XCI during early embryogenesis. We have uncovered a novel function of *XACT* in influencing cell fate decisions during hematopoietic differentiation and in recruitment of *cis*-regulatory proteins and transcriptional regulation. Our research also shows a strong correlation between the expression of *XACT* and the blast origin in AML patients, positioning *XACT* xlncRNA as a potential biomarker to distinguish between various AML subtypes.

## Materials and Methods

### Cell Culture

MEL cells were grown in Dulbecco’s Modified Eagle Medium (DMEM; Gibco, cat. # 41966-029) supplemented with 1% Penicillin-Streptomycin (PS) and 10% fetal bovine serum (FBS). K562 cells were grown in RPMI-1640 medium-PS (Gibco, GlutaMAX supplement, cat. # 61870044) and 10% FBS. HEK293T cells were cultured in DMEM-PS and 15% FBS. All cell cultures were incubated at 37°C, 5% CO_2_, and 95% relative humidity. Adherent cells were maintained at about 80% confluency, while suspension cells were kept at a density of less than 10^6^ cells/ml. All types of cell were split every 2 or 3 days. For the differentiation of K562 cells, cells were seeded at a density of 2.5 x 10^5^ cells/ml and cultured for 72 hours. Hemin (Sigma-Aldrich, cat. # 51280) was dissolved in a solvent containing 0.5M sodium hydroxide and 1M Tris-HCl (pH 8.0), and was used to induce erythroid differentiation at various concentrations of hemin. Different concentrations of phorbol 12-myristate 13-acetate (PMA; Sigma–Aldrich, cat. # P8139) were prepared to induce megakaryocytic differentiation.

HUDEP2 cells were established by others ^81^. This cell line was cultured ^82^ in conditions of 5% CO_2_, 20% O_2_, and 95% ± 5% relative humidity at 37°C. For their expansion, HUDEP2 cells were maintained at a density of 10^6^ cells/ml, and received fresh expansion medium every two-three days. To induce differentiation, the cells were seeded at a density of 2×10^6^ cells/ml in differentiation medium, which was replaced every two-three days, maintaining the density at 2×10^6^ cells/ml. The basic medium primarily consisted of IMDM (PAN-biotech, cat. # P04-20250), supplemented with 1 mg Polyvinyl Alcohol (Sigma-Aldrich, cat. # 363081), 2.5 μg cholesterol/ml (Sigma-Aldrich, cat. # C3045), 2.5 μg L-α-phosphatidylcholine/ml (Sigma-Aldrich, cat. # P3556), 1.5 μg oleic Acid/ml (Sigma-Aldrich, cat. # O1383), 300 μg human holotransferrin/ml (Sanquin, Amsterdam), 10 μg insulin human solution/ml (cat. # I9278, Sigma-Aldrich), 100 μM sodium pyruvate (Gibco, cat. # 113600), 100 U penicillin-streptomycin/ml (Sigma-Aldrich, cat. # P0781), and 2 mM L-Glutamine (Gibco, cat. # 25030081). For the expansion medium, 2 U erythropoietin/ml (EPREX®, Janssen), 2 μg doxycycline/ml (D9891, Sigma-Aldrich), 1 μM Dexamethasone (Sigma-Aldrich, cat. # D4902), and 100 µg human recombinant Stem Cell Factor/L (ITK Diagnostics, cat. # K0921139) were added. For the differentiation medium, 10 U erythropoietin/ml (EPREX®, Janssen), 3.125 U heparin/L (Heparine LEO, Leo Pharma), and 3% (v/v) human plasma (Sanquin, Amsterdam) were added.

### Acute myeloid leukemia (AML) patient samples

Samples of de novo AML patients were collected from the biobanks of the Erasmus MC dept. of Hematology (Rotterdam, The Netherlands) and the University Hospital Regensburg Internal Medicine dept. (Regensburg, Germany). All patients provided written informed consent in accordance with the Declaration of Helsinki and local and national regulations. Mononuclear cells were isolated from bone marrow or peripheral blood as described previously ^83^, and cryopreserved. AML samples obtained by LUMC are described in ^35^.

### SPLiT-seq in K562 and MEL cells

The single-cell RNA sequencing (scRNA-seq) experiment was conducted using the SPLiT-seq method, following the protocol described in ^30^. Initially, MEL cells (mouse) and K562 cells (human) were fixed, permeabilized separately, and then pooled before starting the three rounds of the split-pool barcoding procedure. The pooled cell population was filtered through a 40-μm strainer first, ensuring the generation of single-cell suspensions. The starting number of cells was determined using an automatic cell counter (Countess™ II Automated Cell Counter, Invitrogen), and cells were diluted to 10^6^ cells/ml. The cells were then transferred to the first 96-well barcoding plate, where barcodes were added to the RNA using reverse transcription (RT), and poly-T (20 mer) and random hexamers as primer. These cells were then pooled and distributed into the second 96-well barcoding plate to add the second barcode via ligation. Subsequently, the cells were pooled again, passed through a 40-μm strainer, and redistributed into the third and final 96-well barcoding plate to add the third barcode. This third barcode, which contains the UMI and a biotin at the 3’-end, was added through ligation. Upon completion of the 3-round barcoding protocol, the cells were pooled and counted with an automatic cell counter, and two sub-libraries, each containing 25,000 cells, were generated. The cells were then lysed, and synthesized cDNA was extracted and purified using Dynabeads™ MyOne™ Streptavidin C1 (Invitrogen, cat. # 65002), followed by the generation of the second DNA-strand. The final product was cleaned using KAPA Pure Beads (Roche, cat. # KK8000). The library used for sequencing was generated using the Nextera XT DNA Library Preparation Kit (Illumina, cat. # FC-131-1024). Finally, the sequencing library was subjected to Illumina HiSeq2500 sequencing, generating a total of 159 million paired-end reads (100-bp paired end reads).

### SPLiT-seq data analysis

First the quality of the raw reads was assessed with FastQC ^84^; bad-quality reads were removed using Trimmomatic ^85^. Then, the SPLiT-seq data was de-multiplexed and analyzed using the SPLiT-Seq de-multiplexing pipeline (version 0.1.4) ^86^. The latter was run in merged mode, using the following parameters: minimum 10 reads/cell and maximum 1 error/read. The merged mode takes into consideration that each well of the first barcoding plate contains two barcodes, one corresponding to the poly-T, the other to the random hexamers. The pipeline uses the reverse reads to extract the barcodes and assign the reads to the cells. After the latter, UMI-tools (version 1.0.1) ^87^ was used to extract the UMIs from the reverse reads. The pipeline also uses the selected reverse reads to find their corresponding forward mate paired read. Forward mate paired reads containing the cDNA were aligned to a mixed species (hg version GRCh38 and mg version GRCm38; both downloaded from Ensembl release 99 ^88^ together with their corresponding GTF files (version 99) genome, using STAR (version 2.7.3a) ^89^. Aligned reads were used to determine the number of reads per gene using HTSeq-count (version 0.11.3) ^90^.

### RNA-fluorescence in situ hybridization (RNA-FISH)

#### Slides preparation

The K562 and patient cell suspensions were attached to glass slides using Cytospin3 (Shandon). The cells were diluted to a concentration of 10^6^ cells/ml in DPBS (Dulbecco’s phosphate buffered saline; Gibco, cat. # 14190-094) and 100 µl of cells were spun at 350 rpm with high acceleration for 2 min. Subsequently, the cells were immediately fixed with 3% paraformaldehyde (Electron Microscopy, cat. # 15710) in PBS, for 10 min at 4°C. After fixation, the cells were permeabilized using 0.5% Triton™ X-100 (Sigma, cat. # T8787-250ML) in PBS for 6 min at 4°C. Finally, the cells were washed with ice-cold 70% ethanol. Glass slides were stored in 70% ethanol at -20°C before RNA-FISH.

#### Labeling of the probes

Probes (see Supplementary Table 10) targeting *XIST* (CTD-2200N19), *XACT* (RP11-35D3, BACPAC Genomics) and UTX (RP11-256P2, BACPAC Genomics), respectively, were labeled with fluorescent (fluo) dUTPs using the Nick Translation Kit (Abbott, cat. # 07J00-001) following the manufacturer’s instructions. In brief, 1μg bacterial artificial chromosome (BAC) was mixed with 2.5μl of 0.2mM fluo-dUTP, 5μl of 0.1mM dTTP, 10μl of 0.1mM dNTP mix (GeneON, cat. # 110-012XL) 5μl of 10X nick-translation buffer, 10μl of nick translation enzyme mix (DNase I and E. coli DNA polymerase I) and H_2_O, to a final volume of 50µl. The mixture was then incubated at 16°C overnight, and the reaction stopped by heat inactivation at 70°C for 10 min. The newly generated probes were passed through ProbeQuant G-50 Micro Columns (Cytiva, cat. # 28903408) to remove unused fluo-dUTPs. *XIST* was labeled with Cy5 (Amersham, cat. # PA55022), UTX with red (Abbott, cat # 02N34-050), and *XACT* with green (02N32-050, Abbott).

#### Probe hybridization

Prior to hybridization, a mixture was prepared by combining 3-5μl of labeled probe, 3μl Human Cot-1 DNA (Invitrogen, cat. # 15279-011), 1μl salmon sperm DNA (Invitrogen, cat. # 15632-011), and H_2_O till 50μl. The mixture was gently mixed, and 0.1 volume (5μl) of 3M sodium acetate at pH 4.8 was added and gently mixed, followed by the addition of 125μl of 100% ethanol, and the tube was inverted to mix the contents thoroughly. It was then stored at -20°C for 20 min to precipitate the probes. Subsequently, the probes were centrifuged at maximum speed (13,300 rpm) for 15 min at 4°C. After removing the supernatant, the pelleted material was washed with 1ml of 70% ethanol and centrifuged again, this time for 5 min at 4°C. The ethanol was then removed and the pellet dried for about 10 min. Then, 7μl of 100% formamide was added per spot (Acros Organics, cat. # 327235000) and incubated for 10 min at 75°C for denaturing the probes. The sample was then chilled on ice, and 7μl of cold 2X hybridization buffer per spot was added. The hybridization buffer contained 4XSSC (UltraPure™ SSC, 20X, Invitrogen™, cat. # 15557044), 20% dextran sulfate (BioChemika, cat. # 31403), 2mg BSA/ml (Jena Bioscience, cat. # BU-102) and 40mM Vanadyl Ribonucleoside Complex (New England Biolabs, cat. # S1402S). The mixture was then vortexed, and spun to get rid of bubbles. Before probe hybridization, glass slides were dehydrated by washing for 2 min first with 70% ethanol, followed by 90% ethanol, and finally with 100% ethanol. After these washing steps, slides were air-dried at room temperature. Finally, the probe mix was placed on the dry glass slide and covered with a coverslip. The slides were incubated in 50% formamide in 2XSSC, in a humid chamber, at 37°C overnight.

These slides were then subjected to a series of washing steps. They were first washed twice with 25ml of 50% formamide in 2XSSC (pH 7.2-7.4) for 5 min each. Then, the slides were washed once with 25ml of 2X SCC with Hoechst 33342 (10mg/ml solution in water; Molecular Probes, cat. # H-3570) for 3 min to stain nuclear DNA, and once with 25ml of 2XSCC only, for 2 min. All the washing steps were performed in a water bath at 42°C. Subsequently, the slides were prepared for imaging by mounting them with ProLong Gold anti-fade reagent (Invitrogen, cat. # P36930).

#### Microscope and image analysis

All images were acquired with a Leica Stellaris 5 with HyD S detectors using a 63X oil immersion objective. Z-stacks were collected at 0.5µm steps across the nucleus for the following wavelengths: 405nm (DNA), 488nm (*XACT*), 561 (UTX), and 638nm (*XIST*). Images were acquired with frame average set to 3. Stacks were processed using ImageJ and 2D maximum intensity projections were generated to identify and count the spots.

### RT-qPCR

#### RNA isolation

Total RNA was isolated with ReliaPrep™ RNA Cell Miniprep System (Promega, cat. # Z6012) following the manufacturer’s instructions. In brief, 5×10^5^ to 2×10^6^ cells were lysed with 250µl of BL + TG Buffer by pipetting up and down 7-10 times. Then, 85µl of 100% isopropanol was added, and the sample was mixed by vortexing for 5 seconds. The lysate was transferred to a ReliaPrep™ Minicolumn and centrifuged at 12,000-14,000×g for 30 seconds at room temperature. 500µl of RNA Wash Solution was added, and the column was again washed by centrifugation. An on-column treatment with DNase was performed by adding a mixture of 24µl of Yellow Core Buffer, 3µl 0.09M MnCl_2_, and 3µl of DNaseI enzyme. The sample was incubated for 15 min at room temperature. Then, 200µl of Column Wash Solution was added to the ReliaPrep™ Minicolumn. The sample was centrifuged at 12,000-14,000×g for 15 seconds, 500µl of RNA Wash Solution was added, followed again by centrifugation, now for 30 seconds. Later, 300µl of RNA Wash Solution was added, and centrifuged at 14,000×g for 2 min. RNA was eluted in 30µl nuclease-free water, and collected after centrifugation, now for 1 min, at room temperature.

The isolated RNA was treated with TURBO DNA-free Kit (Ambion, cat. # AM1907) following the manufacturer’s protocol. Briefly, an aliquot of 44µl of RNA (at 200 µg/ml) was mixed on ice with 1µl of Turbo DNase and 5 µl of 10X Turbo DNase Buffer. After incubation at 37°C for 30 minutes, 5µl of RNase Inactivation reagent was added, and the sample incubated for 5 min at room temperature whilst mixing 2-3 times. The reaction was centrifuged at 10,000×g for 90 sec, and the supernatant containing the RNA was transferred to a new tube.

#### cDNA synthesis

RNA was used for cDNA synthesis with SuperScript™ IV Reverse Transcriptase (Invitrogen, cat. # 18090200) and a mix of poly-T and/or random hexamer primers following the manufacturer’s protocol. In brief, 1µg of total isolated RNA was mixed with 1µl of 10mM dNTP mix, 0.25µl of 100µM poly-T, and/or 0.25µl of 100µM random hexamer, and nuclease-free water till a final volume of 13µl. The reaction was incubated at 65°C for 5 min (Biometra TAdvanced PCR thermal cycler, Analitikjena), followed by at least 1 min on ice. Next, 4µl of 5x SuperScript IV Buffer, 1 µL of SuperScript IV Reverse Transcriptase (200U/µl), 1µl of 100mM DTT, and 1µl RNaseOUT Recombinant Ribonuclease Inhibitor (Invitrogen, cat. # 10777019) were added to the annealed RNA. Finally, the reaction was incubated at 23°C for 10 min(when using random hexamer), at 55°C for 10 min and lastly at 80°C for 10 min. The cDNA was diluted as needed with 10mM nuclease-free water.

#### qPCR

PowerUp™ SYBR™ Green Master Mix (Applied Biosystems, cat. # A25741) was used for the qPCR following the manufacturer’s instructions. Briefly, 5µl of 2x PowerUp SYBR Green Master Mix, 0.5µl of 10µM each of forward and reverse primer, 1µl of the cDNA, and 3µl of water were mixed (final volume 10µl). The reaction was done in a CFX96 Touch Real-Time PCR thermal cycler (Bio-Rad) using the following program: 50°C for 2 min, 95°C for 2 min, and 40 cycles of 95°C for 15 sec, 55°C for 15 sec, and 72°C for 1 min. If using Platinum™ Taq DNA-Polymerase, mix cDNA, forward and reverse primer, PCR buffer, MgCl_2_, dNTP mix, DNA Polymerase, SYBR green I nucleic acid gel stain (Sigma-Aldrich, cat. # S9430), we added water to a final volume of 25µl. This mix was subjected to the following program: 94°C for 2 min, and 40 cycles of 94°C for 30 sec, 55°C for 30 sec, and 72°C for 1 min. The melt curve was generated by incubating the reaction at 95°C for 10 sec, followed by 15 sec at each 0.5°C increment between 60°C and 95°C.

### Inactivation of *XACT* using CRISPRi

The plasmid DNA from gRNA-containing vector pBA950 (Addgene, cat. # 122239) was digested by *Bst*XI (NEB, cat. # R0113S) and *Blp*I (NEB, cat. # R0585S). sgRNA targeting *XACT*, or a random gene desert region (non-target, NT), was designed and assembled with the digested gRNA vector using NEBuilder HiFi DNA Assembly kits (NEB, cat. # E2621S). The reaction was then dialyzed using MCE Membrane Filter (Millipore, cat. # VMWP04700), followed by the transformation of competent *E. coli* cells (NEB, cat. # C2987H) using heat shock. *E.coli* cells were also transformed with a plasmid expressing dCas9-Krab (Addgene, cat. # 154473). These two different kinds of transformed bacteria were plated on Ampicillin-agar plates (100 μg Ampicillin/ml) at 37°C overnight, respectively. This was followed by colony picking, individual colonies expanded in LB medium, which contained 20.6mg LB Broth EZMix powder/mL (SIGMA, cat. # L7658) and 100 μg Ampicillin/ml, and plasmid DNA extraction using QIAprep Spin Miniprep Kit (QIAGEN, cat. # 27106) or NucleoBond Xtra Midi kit (Macherey Nagel, cat. # 740410.50) as needed.

HEK293T cells were seeded at a density of 5.8×10^5^ cells/ml in Opti-MEM I (Gibco, cat. # 51985-034)-5% FBS-200 μM sodium pyruvate (Gibco, cat. # 11360070) and incubated overnight. The above two different plasmid DNAs were separately transfected into HEK293T cells, along with three packaging plasmids, i.e. pMDLg/pRRE (Addgene, cat. # 12251), pRSV-Rev (Addgene, cat. # 12253), and pMD2.G (Addgene, cat. #12259), using Lipofectamine™ 3000 Transfection Reagent (Invitrogen, cat. # L3000015). The medium was refreshed 6 hours later, and the generated lentiviruses were harvested 48 hours post-transfection.

HUDEP2 cells were transduced with the lentiviruses containing dCas9-Krab at a multiplicity of infection (MOI) of 0.8. Two days later, cells were transduced again with lentiviruses containing sgRNA. As HUDEP2 cells produce Kusabira Orange, which has spectral overlap with mCherry (dCas9-Krab), they were sorted for BFP (sgRNA) using FACS. Sorted cells were cultured like unprocessed HUDEP2 cells.

### RNA sequencing (RNA-seq) data generation

Total RNA, extracted as described above, was sent to Macrogen for sequencing. The sequencing libraries were constructed using the TruSeq Stranded mRNA LT Sample Prep Kit, following the TruSeq Stranded mRNA Sample Preparation Guide (Part # 15031047). Samples were sequenced on NovaSeqX (150bp pair-end, 30M total reads per sample).

### RNA-seq data analysis

RNA-seq data was generated as described above or obtained from public sources. RNA-seq datasets of two AML patient cohorts were obtained from the European genome-phenome archive (EGA) after permission was obtained from the data generators (with acc. no. EGA S00001004684 for LUMC ^35^, and EGA C00001000956 for EMC ^36^). BAM files and read counts/gene were obtained from EGA and/or directly from the authors. BAM files were used for visualization of the RNA-seq data. For other datasets, the raw sequencing data from published RNA-seq dataset was acquired using Kingfisher v0.1.2 ^91^. Low-quality bases and adapters were trimmed by Fastp v0.23.2 ^92^. The quality of both the raw reads and cleaned reads was examined by FastQC v0.11.9 ^84^. The cleaned reads were then aligned to the GRCh38 reference genome using Hisat2 v2.2.1 ^93^, with the ‘new-summary’ parameter, followed by BAM file sorting, indexing using Samtools v1.6 ^94^. Optionally, ‘Samtools view’ was used to obtain BAM files containing only chromosome X information. Bigwig files were generated from the sorted BAM files with bamCoverage – deeptools v3.5.1 ^95^. Sequencing read counts of dataset were obtained using htseq-count v2.0.2 ^90^, or featureCounts v2.0.3 ^96^. Normalized counts and differentially expressed genes were obtained using DESeq2 package v1.38.0 ^97^. ComBat-seq ^98^ was used to adjust batch effect when necessary. The normalized counts were used for downstream processing such as plotting X:A ratio, plotting gene correlation, chromosome wide gene expression profiling, and identification of differently expressed genes. In addition, the regions of interest were visualized through Integrative Genomics Viewer (IGV) v2.12.3 ^99^ or UCSC genome browser ^100^. Enrichr ^101^ and g:Profiler ^102^ were used to perform enrichment analysis of the genes of interest. A cutoff of 0.05, based on the Benjamini-Hochberg FDR method was applied for g:Profiler. CIBERSORTx ^53^ was used to evaluate the abundance of particular cell types among samples. Protein-protein association networks were investigated using STRING ^103^, and the results were visualized with Cytoscape ^104^. Important modules and hub nodes were identified using MCODE ^54^and Cytohubba ^55^, respectively, which are applications available in Cytoscape.

### Cut&Tag data generation

Cut&Tag experiments were performed with CUT&TAG Assay kit (Active Motif, cat. # 53160). Briefly, 5×10^5^ fresh cells per sample were harvested and washed with wash buffer, then bound to pre-washed Concanavalin A beads at room temperature for 10 min. Next, the samples were incubated with corresponding primary antibody in antibody buffer at 4°C overnight. The primary antibodies included anti-H3K27ac (Active Motif, cat. # 39133), anti-H3K27me3(Active Motif, cat. # 39155), and Rabbit Anti-Mouse IgG (Abcam, cat. # ab46540). The liquid was removed the next day, and samples were incubated with guinea pig anti-rabbit antibody at room temperature for 1 hour. This was followed by three washes with Dig-wash buffer, and incubation with assembled pA-Tn5. Following incubation, the samples were washed three times with Dig-300 buffer. The samples were then incubated at 37°C for 1 hour with Tagmentation Buffer. Tagmentation was stopped by adding EDTA (final concentration of 0.0168M) EDTA, as well as (final) 0.1% SDS and 0.085mg proteinase-K/ml. Tagmented DNA was extracted using the column-based method, as per the protocol. Tagmented DNA was PCR amplified and prepared for sequencing by the Erasmus MC Biomics-Genomics core facility. The libraries were sequenced on an Illumina NextSeq2000 sequencer. Paired-end clusters were generated, of 50 bases in length.

### ATAC-seq data generation

Live cells (50,000 cells per sample) were harvested, washed twice with PBS-5% FBS. Sequencing library was prepared using OMNI-ATAC ^105^ protocol. These libraries were subsequently sequenced on an Illumina Nextseq2000 sequencer. Paired-end clusters of 50 bases in length were generated and sequenced.

### ATAC-seq and Cut&Tag data analysis

Raw sequencing data was trimmed using Fastp ^92^ and examined using FastQC ^84^. Sequencing reads that fulfilled quality requirements were aligned to the human GRCh38 genome using Bowtie2 ^106^, followed by BAM file viewing, sorting, and indexing using Samtools ^94^. This was followed by Picard ^107^ and ‘bedtools intersect’ ^108^ to remove duplicate reads and blacklisted regions. MACS3 ^109^ was used to call peaks (narrowPeaks for ATAC-seq (qvalue = 0.001) and H3K27ac (pvalue = 0.01), and broadPeaks for H3K27me3 (pvalue = 0.01)).

Specifically, for histone modification samples, IgG sample from HUDEP2-NT served as a control for the corresponding HUDEP2-NT samples, while IgG sample from HUDEP2-KD was used as a control for those samples from HUDEP2-KD cells. Raw counts of peaks were obtained with Diffbind ^110^. Next, differentially enriched peaks (DE peaks) were identified with edgeR ^111^. The cutoffs for ATAC-seq and H3K27me3 are FDR <= 0.05 and |log2FC| > 1, while the cutoffs for H3K27ac are FDR <= 0.05 and |log2FC| > 0.5. ComputeMatrix (deepTools) ^95^ was used to assess scores for specific genomic regions, which were then visualized through plotHeatmap (deepTools). The observation of genomic feature distribution of DE peaks was performed through ChIPseeker^112^. GREAT ^113^ was employed to annotate DE peaks.

#### Motif analysis

Motif discovery was performed using Homer ^114^ on both highly and lowly enriched narrow peaks. For H3K27me3, notably, DE peaks covering TSS were extracted, which were then extended 200bp upstream and downstream of the TSS for motif analysis. The significant motifs (p-value <= 0.01) from highly and lowly enriched peaks were merged using the ‘full_join’ function in the Tidyverse package^115^. Missing values of log_Pvalue were assigned a value of “-1”. To reduce background noise, motifs were ranked by the ratio of “log_Pvalue(highly enriched) / log_Pvalue(lowly enriched)” and the ratio of “log_Pvalue(lowly enriched) / log_Pvalue(highly enriched)”, so that the most significant motifs could be distinguished in the HUDEP2-KD and HUDEP2-NT samples, respectively.

#### Footprint analysis

For ATAC-seq and H3K27ac data, bam files from replicates in the same group were merged by ‘samtools merge’ function, while the reproducible narrow peaks for each group were obtained with the use of IDR ^116^. Footprints were then identified with TOBIAS ^59^. ‘TOBIAS ATACorrect’, ‘TOBIAS ScoreBigwig’, and ‘TOBIAS BINDetect’ were performed in sequence to correct bias, calculate footprint scores, and estimate differentially bound motifs.

### Function enrichment analysis and visualization

Gene ontology (GO) enrichment analysis and visualization were conducted according to the protocol established by *Reimand et al. ^117^*. The GO terms in the biological process (BP) and molecular function (MF) categories were analyzed using g:Profiler ^102^, applying the Benjamini-Hochberg FDR method. For RNA-seq, ATAC-seq, and H3K27me3, a cutoff of 0.05 was used, while the cutoff for H3K27ac was 0.1. The functional enrichment analysis results were visualized using EnrichmentMap ^118^, with a node cutoff (Q-value) of 0.01, and an edge cutoff (Similarity) of 0.6. Subsequently, AutoAnnotate ^119^ was used to identify clusters of similar nodes. The labels for these clusters were based on the enrichment results from RNA-seq with the minimum FDR Q-value.

## Data availability

Datasets that were generated in this study are available at the NCBI GEO database with accession numbers (ATAC-seq, GSE283510; Cut&Tag, GSE283512; RNA-seq, GSE283513)

The publicly available data used in this study can be found in the following online repositories. RNA-seq datasets of AML patients are available from the European Genome-phenome Archive (EGA): LUMC dataset (accession number EGA C00001000956) and EMC dataset (accession number EGAS00001004684).

## Acknowledgements

We thank the China Scholarship Council for financially supporting P.Z., H.Z., and T.L. We also extend our gratitude to Rıdvan Cetin (Department of Cell Biology, Erasmus MC) and Joana Carvalho Mereira de Mello (Department of Developmental Biology, Erasmus MC) for their valuable advice on designing the CRISPR knockdown and FISH experiments, respectively.

## Author contributions

E.M. conceived the original idea, supervised the project, analyzed and interpreted the data, and wrote the manuscript. P.Z. conducted experiments, analyzed and interpreted the data, and wrote the manuscript with E.M.. P.H.A. conducted parts of the experiments, participated in data analysis, and contributed to manuscript writing. H.Z. assisted in experimental design, performed data analysis, and contributed to manuscript writing. C.R. contributed to data analysis, interpretation, and manuscript editing. D.H. participated in data interpretation, and contributed to manuscript writing. F.G., participated in data interpretation, and contributed to manuscript editing. J.G. contributed to FISH experiments, and manuscript editing. W.IJ. Performed library preparation for ATAC-seq and Cut&Tag, conducted initial data analysis, and contributed to manuscript editing. T.L. and S.J. assisted with HUDEP2 knockdown experiments and data interpretation. R.M.L. and R.D. assisted with AML patient data acquisition, data analysis and interpretation, and provided AML patient samples, and contributed to manuscript editing. All authors read and approved the final manuscript.

## Ethics declaration

All authors declare that they have no conflicts of interest.

## Supplementary Figures legends

**Supplementary Figure 1. Detailed description of the *XACT* locus.**

**A)** The region shown is taken from human genome GRCh38: chrX:113,458,727-114,143,714); the different annotated transcripts are in green. The originally described 252 kb-long *XACT* transcript (as reported in Vallot et al., 2013) is displayed in pink. The *T113.3* gene is located downstream of *XACT* and overlaps with the longest *XACT* transcript we mapped. The suggested transcription start sites (TSS) of *XACT* and *T113.3*, respectively, are marked by vertical red lines. **B)** *XACT* expression in publicly available bulk RNA-seq data generated from K562 cells (Dataset: ERR688). The region shown is human genome GRCh38/hg38: chrX:113,458,727-114,143,714). The light blue area represents the locus encompassing *XACT*. K562-R1 and -R2 represent replicated experiments. The y-axis indicates the number of counts (range was set to 100). Data was visualized using Integrative Genomics Viewer (IGV).

**Supplementary Figure 2. Histone marks and expression profiles within the locus encompassing *XACT* in K562 cells, human embryonic stem cells (hESCs), and human induced pluripotent cells (hiPSCs).**

**A)** *XACT* expression in K562 cells, as generated by SPLiT-seq based RNA-seq (see also Figure 1A). The total RNA-seq count of all cells is displayed on the y-axis. The region shown is GRCh38/hg38: chrX:113,458,727-114,143,714. The light blue area represents the locus encompassing *XACT*. **B)** H3K27ac ChIP-seq data at the *XACT* locus in K562 cells, H1 and H9 hESCs, and hiPSCs. The H3K27ac ChIP-seq data from ENCODE is visualized using the UCSC genome browser. Pooled fold-change of ChIP-seq experiments (from replicates) over control is depicted on the y-axis. **C)** H3K4me3 ChIP-seq data at the *XACT* locus in the same sets of cells. H3K4me3 ChIP-seq data is shown, using an identical approach as in B.

**Supplementary Figure 3. RNA-seq read counts distribution, and validation by RT-qPCR of *XACT* expression in AML patients.**

**A)** RNA-seq read counts distribution plot in AML patient samples. Reads were extracted from the *XACT* locus and their counts were plotted in each patient (left: EMC data from 150 patients, right: LUMC data from 100 patients). **B)** *XACT* transcript reads (RNA-seq) in selected patients (indicated as male and female) of the EMC AML dataset. Reads that map to the minus strand (see main text) are displayed in red, those that map to the plus strand are in blue. **C)** RT-qPCR results of selected patients with the operationally defined high vs. low expression of *XACT*. The delta CT-value (dCTq) is displayed on the y-axis (low dCTq = high expression, and high dCTq = low expression).

**Supplementary Figure 4. X*A*CT expression correlation with *XIST*, and impact on dosage compensation (for the LUMC cohort data).**

**A)** *XACT* and *XIST* levels (as log2 counts per million, cpm) are depicted on the x- and y-axis, respectively, separately for males and females. Male patients are plotted in turquoise, females in reddish-orange. **B)** X:A (X-chromosome to autosome) ratio of the LUMC AML patient cohort. The *XACT* level in each patient is defined as high (circle) or low (triangle), and this both for males (turquoise) and females (orange); the average for each sex is indicated by dashed lines. **C, D)** Boxplots showing average expression of X-linked genes in *XACT-*high (in red) and *XACT*-low (blue) AML patients. The two boxplots in panels C-D also show the average expression in male and female patients, respectively. **E, F)** Correlation of the average expression of X-linked genes in *XACT*-high and *XACT*-low AML patients, for males (in E) and females (F). Genes that show a significant change between *XACT*-high and *XACT*-low (p value <=0.05, T-test) are in red and green, respectively. Green colored dots (together with gene names) show genes with significant change (p value <=0.05 and a |log2FC| >=1.5). The average expression level in *XACT*-high and low patients is shown on the x- and y-axis, respectively. **G, H)** Changes in average expression of X-linked genes along the X-chromosome. Average expression of each gene in *XACT*-high and *XACT*-low AML patients is used to calculate the ratio (as log2 ratio) of each gene depicted on the y-axis. The location of the gene on the X-chromosome is shown on the x-axis and given in mega basepairs (mb). The location of *XACT* is shown with a blue vertical dashed line. Males are shown in G, females in H. The color of each gene is similar to the color used in E, F.

**Supplementary Figure 5. Average expression of chromosome (chr)-3 and chr-8 linked genes across *XACT*-high and *XACT*-low patients (for the LUMC cohort data).**

**A,B)** Boxplots of average expression of chr3-linked genes in male (in A) and female AML patients (B). **C,D)** Boxplots of average expression of chr8-linked genes in male (in C) and female AML patients (D).

**Supplementary Figure 6. Expression and chromatin dynamics of the *XACT* locus during normal hematopoiesis.**

**A)** *XACT* RNA reads across the *XACT* locus in different cell states/subtypes. For each cell state the IGV expression from different donors is overlaid (IGV, Integrative Genomics Viewer), F=female (red), M=male (green). **B)** Chromatin dynamics (using ATAC-seq) of the locus encompassing *XACT*, during normal hematopoiesis, given for different cell subpopulations/states.

**Supplementary Figure 7.**

**A)** X-chromosome to autosome (X:A) ratio in different states of hematopoietic cell differentiation, in male (green) and female (red) donors. **B)** *XACT* expression in different hematopoietic cell states (the publicly available RNA-seq dataset NCBI GSE113182 was used).

**Supplementary Figure 8. Expression and chromatin dynamics of *XACT* in HUDEP2 cells, in proliferative state and after differentiation.**

**A)** Expression of *XACT* in HUDEP2-NT and HUDEP2-KD cells (for the derivation and use of these cells, see Materials and Methods). **B)** *XACT* expression during HUDEP2 differentiation up to day 8. **C)** RNA yield after 3 days of differentiation in HUDEP2-KD and HUDEP2-NT cells, starting with the same number of seed cells. **D-F)** Volcano plots showing the differential chromatin accessibility (D), H3K27ac (E) and H3K27me3 (F) profiles, respectively. **G)** Chromosomal location of the regions identified as DEGs (in RNA-seq), differentially accessible regions from ATAC-seq, and differential profiles of H3K27ac and H3K27me3 marks in HUDEP2-KD, compared to HUDEP2-NT.

**Supplementary Figure 9. Pathway enrichment analysis, and zoom-in.**

Pathway enrichment analysis combining multi-omics data for Group 1 (genes that show downregulation (RNA-seq), regions that show low chromatin accessibility (ATAC-seq), low H3K27ac, and high H3K27me3 (see Supplementary Note1 for grouping) in HUDEP2-KD cells. The different pathways identified in this analysis are depicted on the left. The most significantly enriched cluster (DNA-binding and transcription factor activity, named after the most significant node in this cluster in RNA-seq) is depicted on the right. Each node carries information about the types of data and significance (in colour code). The connections between nodes (edges) are in blue.

**Supplementary Figure 10. Pathway enrichment analysis, and zoom-in.**

Pathway enrichment analysis combining multi-omics data for Group 2 (genes that show upregulation (RNA-seq), regions that show high chromatin accessibility (ATAC-seq), high H3K27ac, and low H3K27me3) (see Supplementary Note1 for grouping) in HUDEP2-KD cells. The different pathways identified in this analysis are depicted on the left. The most two significantly enriched clusters (response to stimulus and anatomical structure morphogenesis, named after the most significant node in each cluster in RNA-seq) are re-clustered and depicted on the right. Each node carries information about the type of data and significance (in colour code). The connections between nodes (edges) are in blue.

## Supplementary Tables descriptions

**Supplementary Table 1. Pooled gene count after SPLiT-seq single-cell RNA-seq experiment.**

Top-5 genes in human (K562) and mouse (MEL) erythroleukemic cells. After scRNA-seq, the total count of each gene in all cells was used for ranking.

**Supplementary Table 2. Enriched gene ontology (GO) driver terms for differentially expressed genes (DEGs) between proliferating HUDEP2-NT and HUDEP2-KD cells.**

DEGs: List of up and down regulated genes in HUDEP2-KD cells.

Down_genes_GO_driver_terms: Biological process and molecular function GO driver terms enriched for genes that are down regulated in HUDEP2-KD cells.

Up_genes_GO_driver_terms: Biological process and molecular function GO driver terms enriched for genes that are up regulated in HUDEP2-KD cells.

**Supplementary Table 3. Enriched gene ontology (GO) driver terms for differentially expressed genes (DEGs) between HUDEP2-NT and HUDEP2-KD after 3 days of differentiation.**

DEGs: List of up and down regulated genes in HUDEP2-KD cells.

Down_genes_GO_driver_terms: Biological process and molecular function GO driver terms enriched for genes that are down regulated in HUDEP2-KD cells.

Up_genes_GO_driver_terms: Biological process and molecular function GO driver terms enriched for genes that are up regulated in HUDEP2-KD cells.

**Supplementary Table 4. Protein-protein interaction analysis, MCODE, and hub genes for differentially expressed genes between HUDEP2-NT and HUDEP2-KD at proliferative state.**

Up_PPI: Short tabular text of protein-protein interaction network based on up-regulated genes in HUDEP2-KD cells(STRING).

Up_MCODE: Molecular complex detection results based on up-regulated genes in HUDEP2-KD cells.

Up_hubgenes: Exploration results of hub nodes based on up-regulated genes in HUDEP2-KD cells.

Down_PPI: Short tabular text of protein-protein interaction networks based on down-regulated genes in HUDEP2-KD cells(STRING).

Down_MCODE: Molecular complex detection results based on down-regulated genes in HUDEP2-KD cells.

Down_hubgenes: Exploration results of hub nodes based on down-regulated genes in HUDEP2-KD cells.

**Supplementary Table 5. Differential accessible regions (DARs) obtained from ATAC-seq between proliferating HUDEP2-KD and HUDEP2-NT cells.**

DARs_genes: List of differential accessible regions (|log2FC|>=1, FDR <=0.05) between HUDEP2-KD and HUDEP2-NT cells, as well as their associated genes annotated by GREAT.

Footprints: Assessment of differential binding motifs between HUDEP2-KD and HUDEP2-NT using TOBIAS.

Motif_ranking: Ratio of the log p-values from known motif enrichment results based on the DARs between HUDEP2-KD and HUDEP2-NT, along with their ranking.

Up_HUDEP2-KD_knownResults: Known motif enrichment results from Homer based on the DARs with higher accessibility in HUDEP2-KD cells.

Down_HUDEP2-KD_knownResults: Known motif enrichment results from Homer based on the DARs with lower accessibility in HUDEP2-KD cells.

**Supplementary Table 6. Differential histone marks of H3K27ac between proliferating HUDEP2-KD and HUDEP2-NT cells.**

DBRs_genes: List of differential binding regions (|log2FC|>=0.5, FDR <=0.05) of H3K27ac between HUDEP2-KD and HUDEP2-NT cells, as well as their associated genes annotated by GREAT.

Footprints: Assessment of differential binding motifs between HUDEP2-KD and HUDEP2-NT using TOBIAS.

Motif_ranking: Ratio of the log p-values from known motif enrichment results based on the DBRs of H3K27ac between HUDEP2-KD and HUDEP2-NT, along with their ranking.

Up_HUDEP2-KD_knownResults: Known motif enrichment results from Homer based on the DBRs of H3K27ac with increased binding ability in HUDEP2-KD cells.

Down_HUDEP2-KD_knownResults: Known motif enrichment results from Homer based on the DBRs of H3K27ac with decreased binding ability in HUDEP2-KD cells.

**Supplementary Table 7. Differential histone marks of H3K27me3 between proliferating HUDEP2-KD and HUDEP2-NT cells.**

DBRs_genes: List of differential binding regions (|log2FC|>=1, FDR <=0.05) of H3K27me3 between HUDEP2-KD and HUDEP2-NT cells, as well as their associated genes annotated by GREAT.

Up_HUDEP2-KD_cover_TSS: Higher H3K27me3 modification regions that cover TSS in HUDEP2-KD cells.

Down_HUDEP2-KD_cover_TSS: Lower H3K27me3 modification regions that cover TSS in HUDEP2-KD cells.

Motif_ranking: Ratio of the log p-values from known motif enrichment results based on the -/+ 200bp of TSS, which are covered by the DBRs of H3K27me3 between HUDEP2-KD and HUDEP2-NT, along with their ranking.

Up_HUDEP2-KD_knownResults: Known motif enrichment results from Homer based on the -/+ 200bp of TSS, which are covered by the DBRs of H3K27me3 with increased binding ability in HUDEP2-KD cells.

Down_HUDEP2-KD_knownResults: Known motif enrichment results from Homer based on the -/+ 200bp of TSS, which are covered by the DBRs of H3K27me3 with decreased binding ability in HUDEP2-KD cells.

**Supplementary Table 8. Enriched gene ontology (GO) terms from multi-omics datasets between proliferating HUDEP2-NT and HUDEP2-KD cells.**

Up in RNA-seq: Biological process and molecular function GO terms enriched for genes up regulated in HUDEP2-KD cells(FDR <= 0.05).

Down in RNA-seq: Biological process and molecular function GO terms enriched for genes down regulated in HUDEP2-KD cells(FDR <= 0.05).

Up in ATAC-seq: Biological process and molecular function GO terms enriched for genes annotated from the highly accessible regions in HUDEP2-KD cells(FDR <= 0.05).

Down in ATAC-seq: Biological process and molecular function GO terms enriched for genes annotated from the lowly accessible regions in HUDEP2-KD cells(FDR <= 0.05).

Up in H3K27ac: Biological process and molecular function GO terms enriched for genes annotated from the regions with high H3K27ac binding in HUDEP2-KD cells(FDR <= 0.1).

Down in H3K27ac: Biological process and molecular function GO terms enriched for genes annotated from the regions with low H3K27ac binding in HUDEP2-KD cells(FDR <= 0.1).

Up in H3K27me3: Biological process and molecular function GO terms enriched for genes annotated from the regions with high H3K27me3 binding in HUDEP2-KD cells(FDR <= 0.05).

Down in H3K27me3: Biological process and molecular function GO terms enriched for genes annotated from the regions with low H3K27me3 binding in HUDEP2-KD cells(FDR <= 0.05).

**Supplementary Table 9. Genes down-regulated in *XACT*-high patients compared to patients with low *XACT* (refer to Figure 6A,B for the overlapping genes between datasets on Venn diagrams).**

Down-regulated in XACT-high: List of down-regulated genes in *XACT*-high AML patients.

Up-regulated in XACT-high: List of up-regulated genes in *XACT*-high AML patients.

Down-regulated in *XACT*-high_GO: enriched GO terms for down-regulated genes in *XACT*-high AML patients.

Up-regulated in *XACT*-high_GO: enriched GO terms for up-regulated genes in *XACT*-high AML patients.

**Supplementary Table 10. Information of RNA-FISH probes, RT-qPCR primers, and sgRNAs for CRISPRi.**

