## Supplementary Figure_1 for "A Novel *XACT* lncRNA Transcript with Functions Transcending X-Chromosome Inactivation"

**A)**

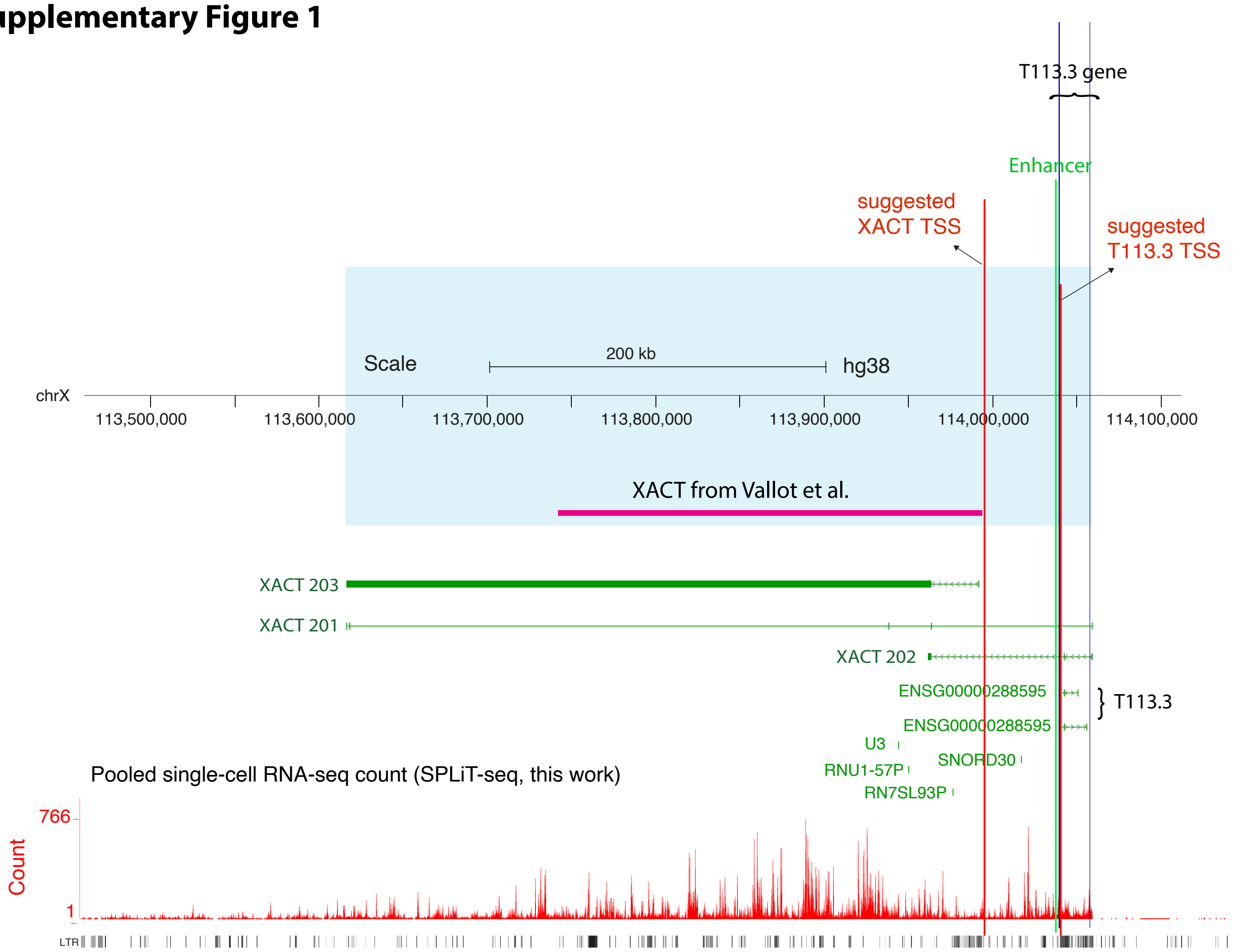


XACT from Vallot et al. (Nat genetics 2013)
Hg19: chrX:112,983,323 -113,235,148
GRCh38: chrX:113,740,044 -113,991,913

XACT -TSS is suggested to be around: GRCh38 chrX:113,995,000

Ensembl:
ENST00000674361.1, XACT-203, hg38 chrX:113,616,300-113,991,976 ( closer to Vallot et al. Nat. genetics 2013 )
ENST00000468762.3, XACT-201, hg38 chrX:113,616,300-114,059,121 ( longest, it also covers T113, but expressed from - strand)
ENST00000649506.1, XACT-202, hg38 chrX:113,961,711-114,059,289 ( shortest, it also covers T113, but expressed from - strand)

T113.3
T113.3 promoter in hg19: chrX:113,283,559 and in hg38: chrX:114,040,388

ESC specific Enhancer
chrX:114,037,565-114,038,230

**B)**

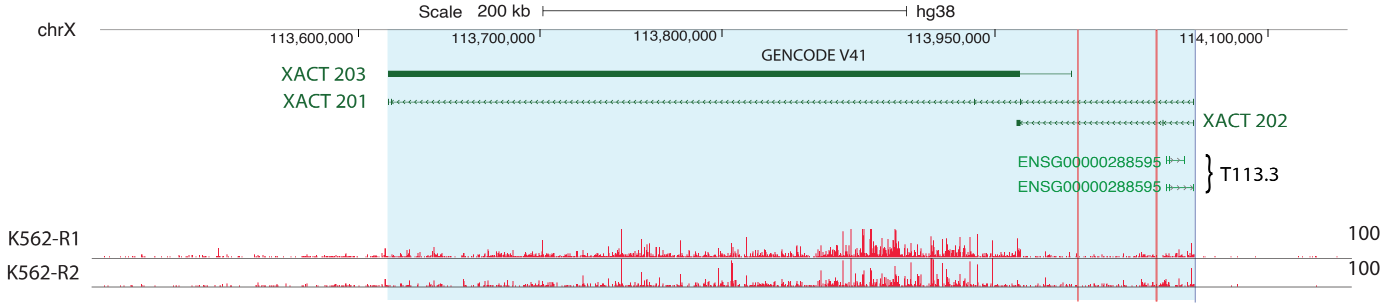


ERR688 dataset, bulk RNA-seq (CeMM2_1, CeMM2_2)

Zhang et al. 2025
