## Supplementary figures and images for "A Novel *XACT* lncRNA Transcript with Functions Transcending X-Chromosome Inactivation"

### Supplementary Figure_2

#### Supplementary Figure 2

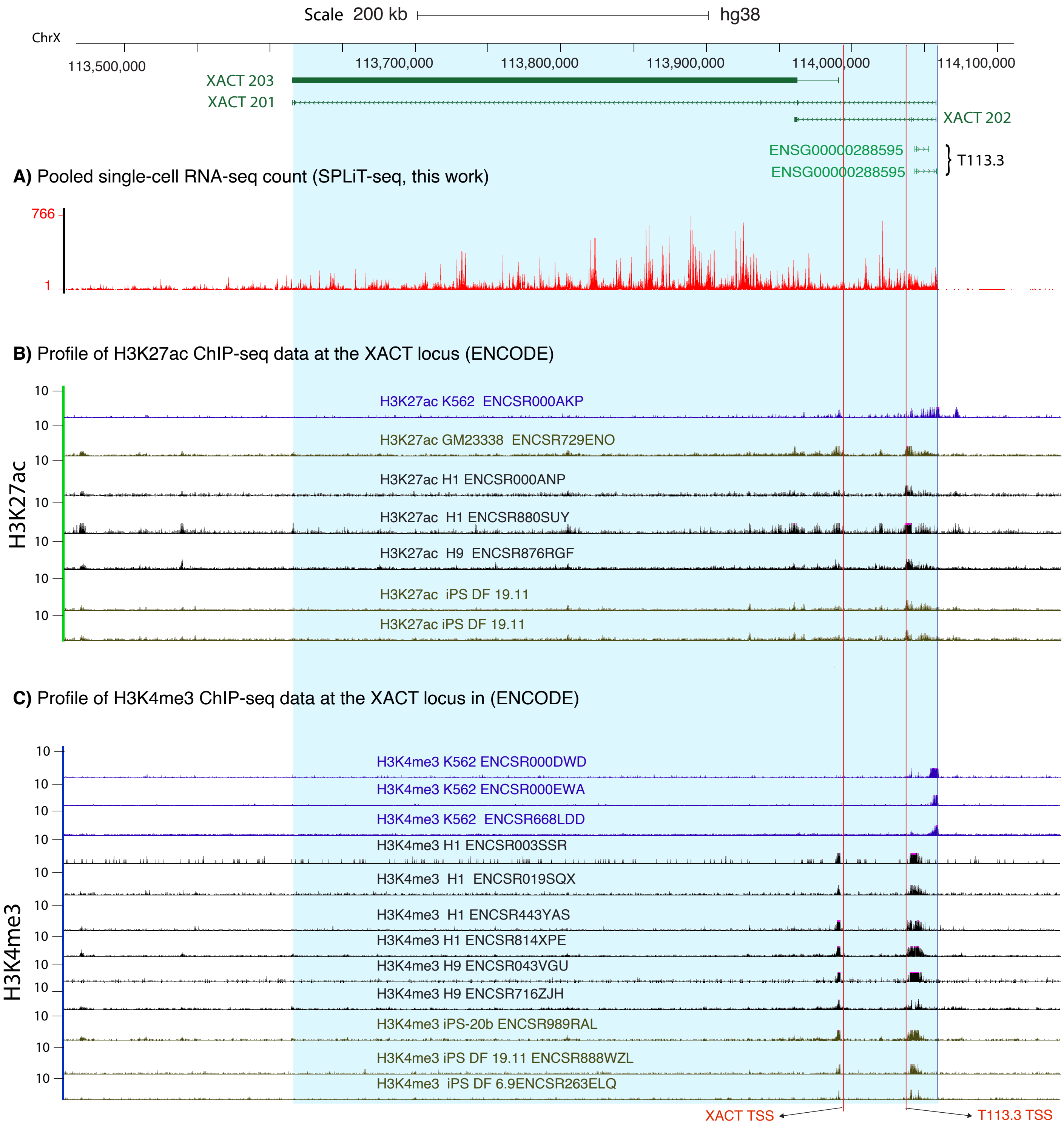


Zhang et al. 2025

### Supplementary Figure_3_C

##### Supplementary Figure 3

**C)** XACT expression in selected patients (RT-qPCR)

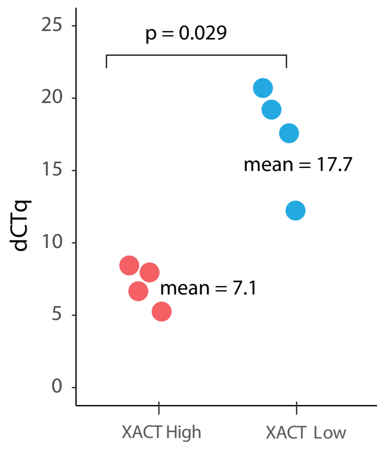

### Supplementary Figure_4

#### Supplementary Figure 4

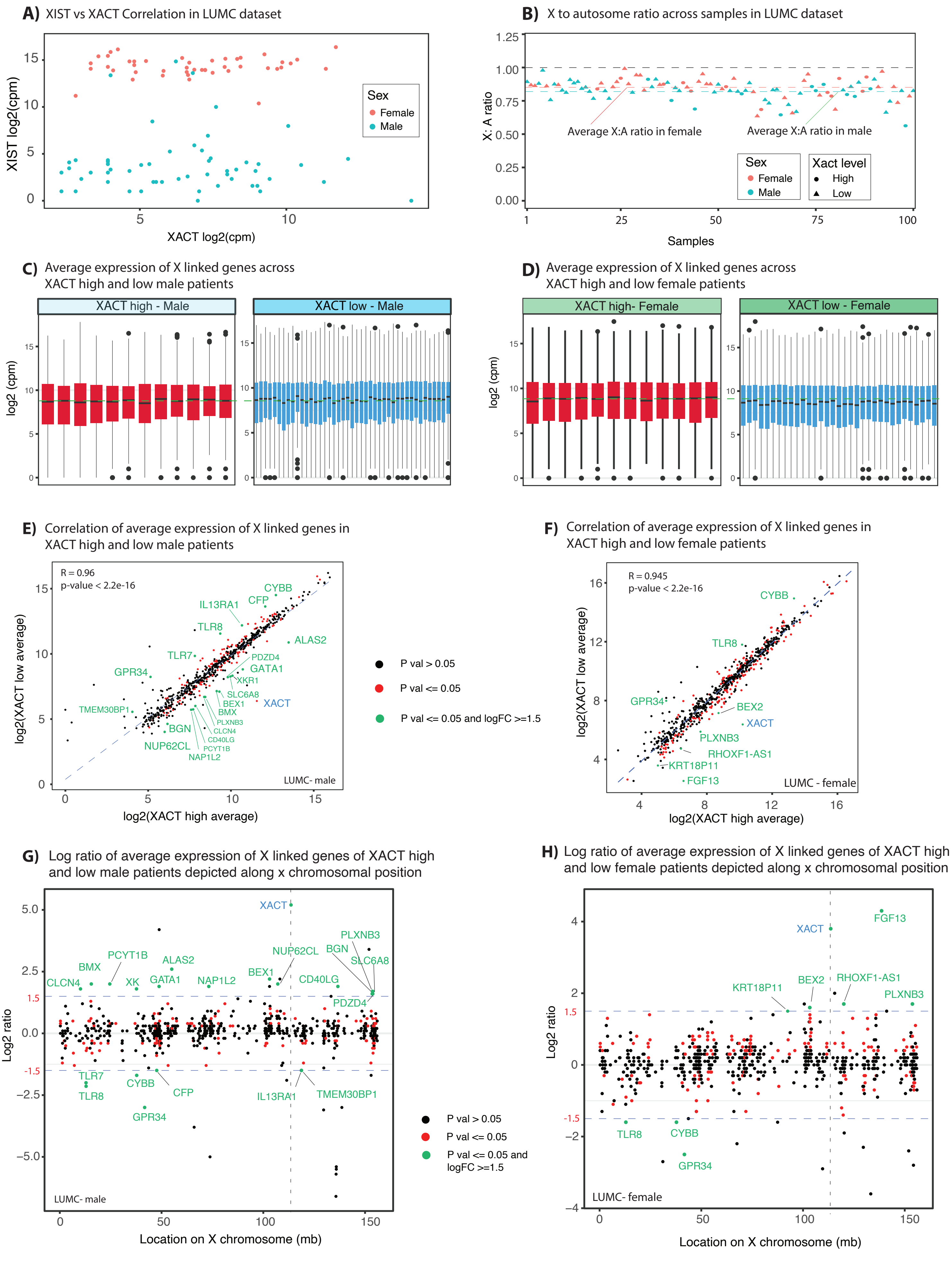


Zhang et al. 2025

### Supplementary Figure_8

#### Supplementary Figure 8

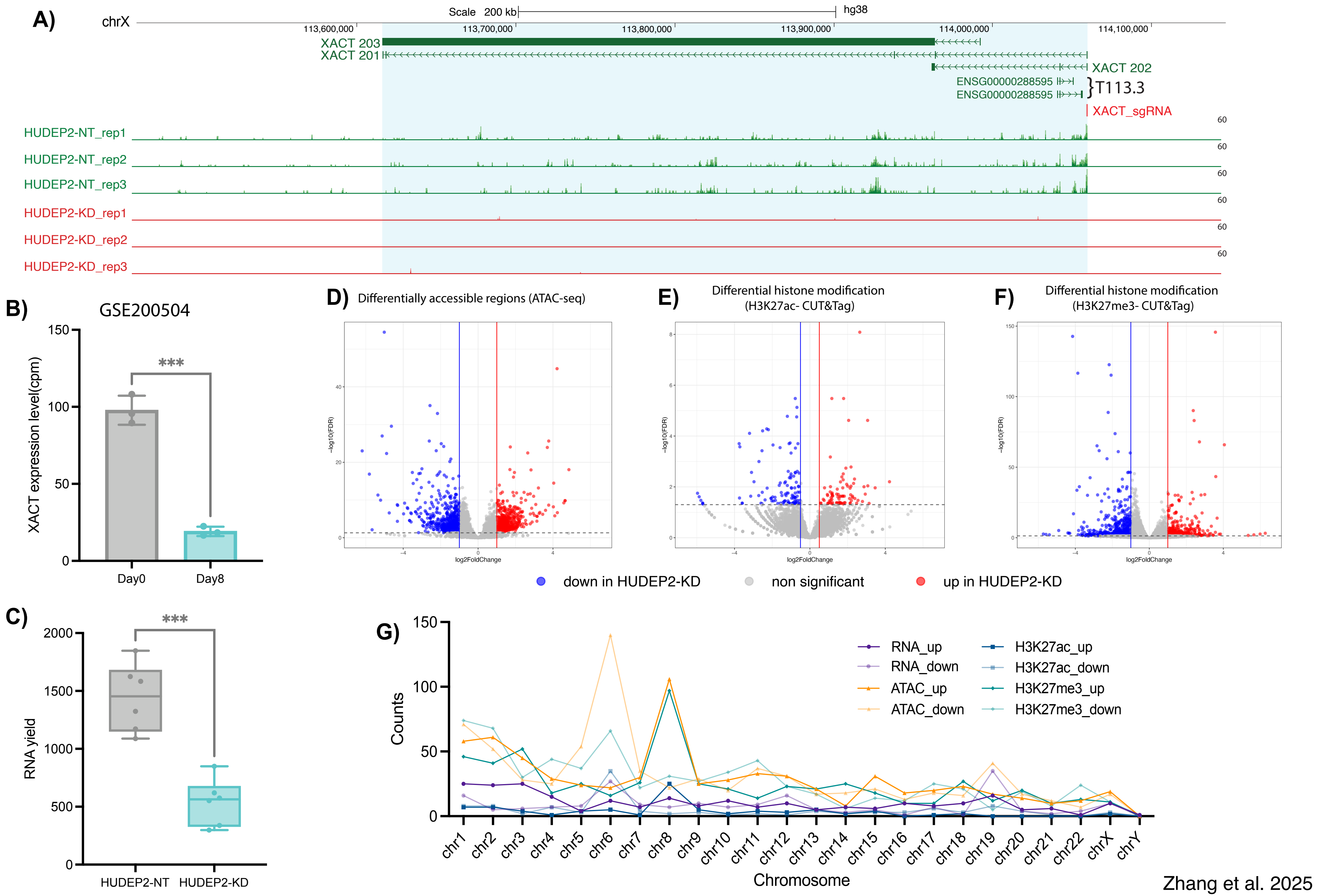


Zhang et al. 2025

### Supplementary Figure_9

##### Supplementary Figure 9

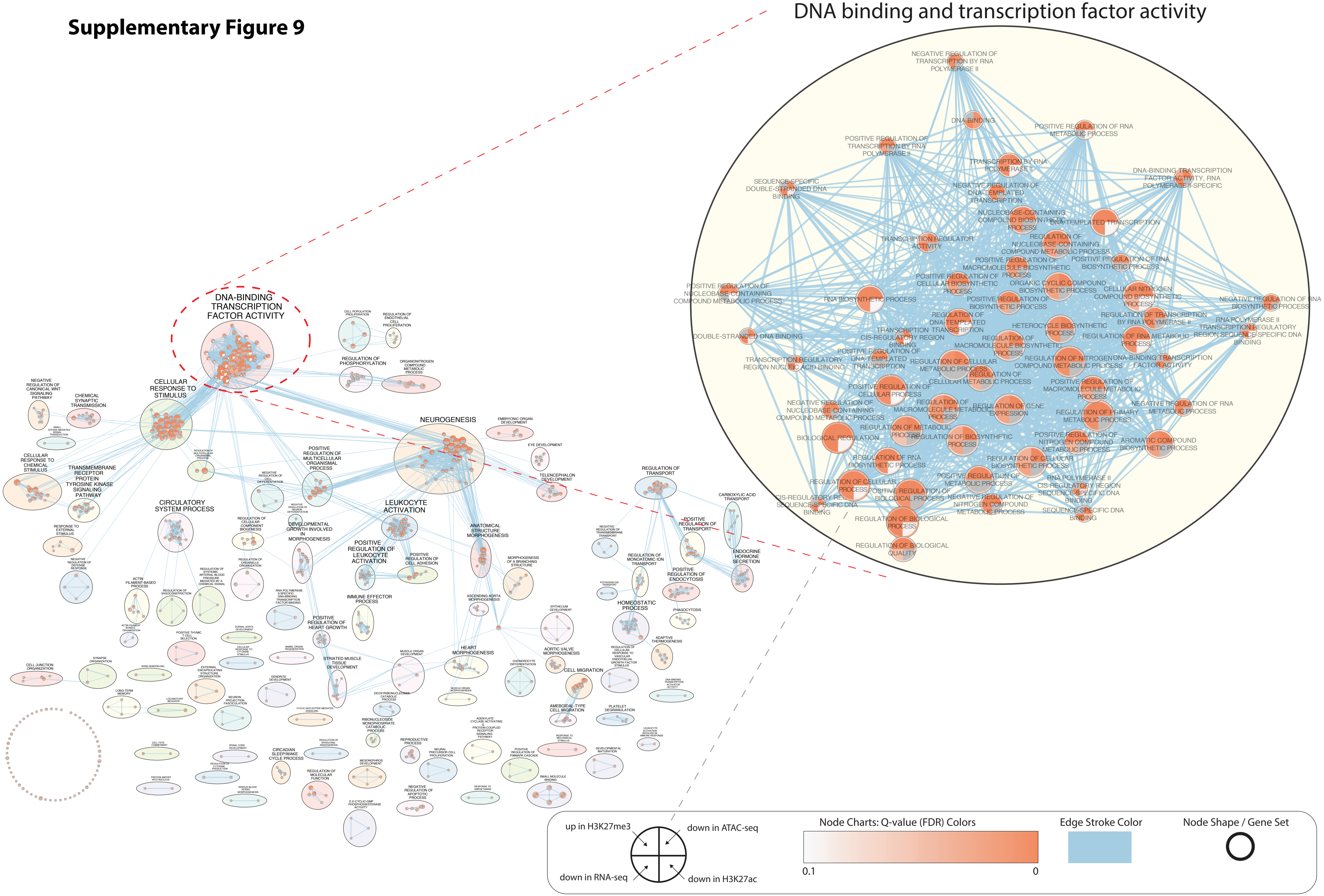


Zhang et al. 2025

### Supplementary Figure_10

**Supplementary Figure 10**

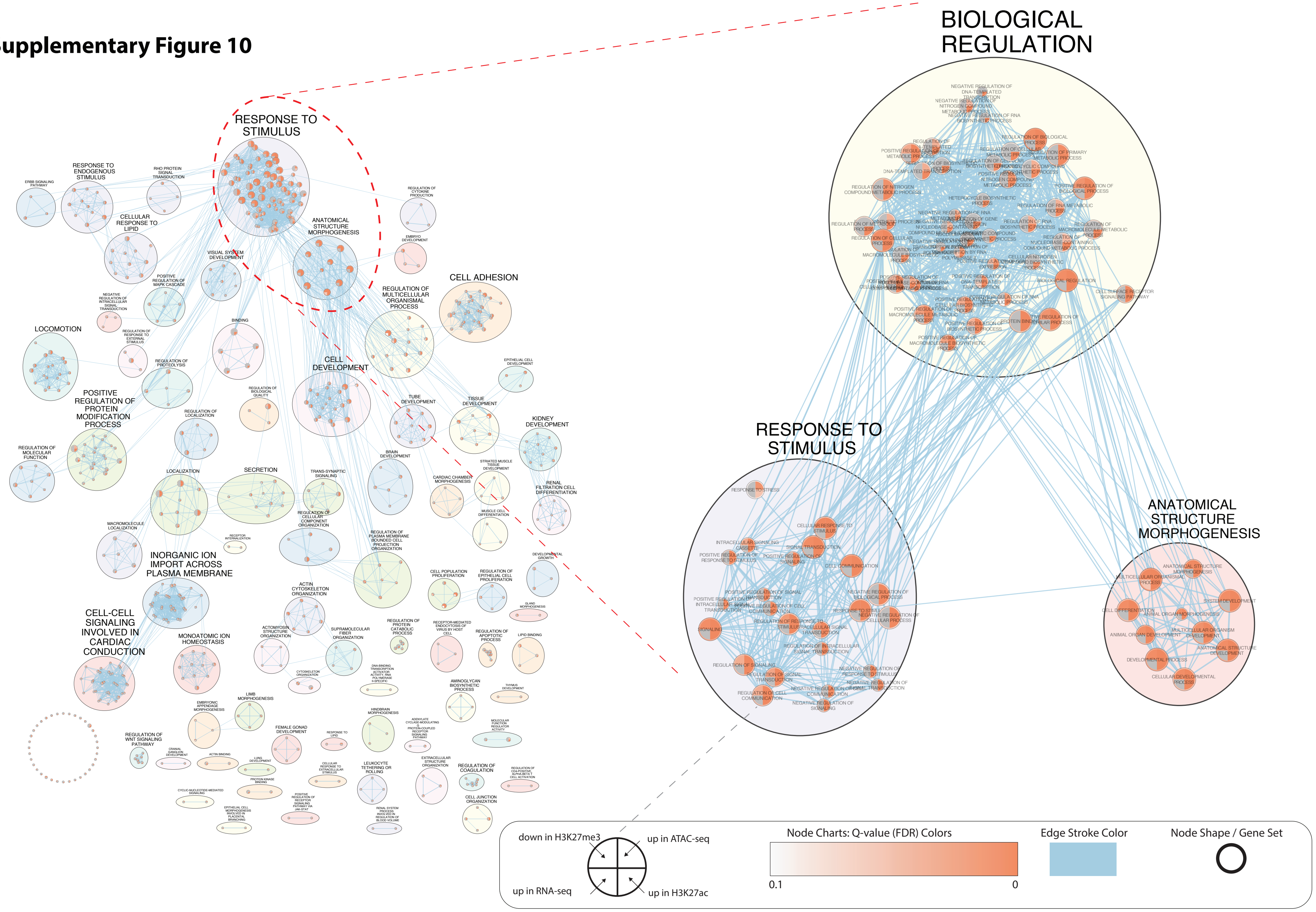


Zhang et al. 2025
