## Supplementary Figure_3_A_B for "A Novel *XACT* lncRNA Transcript with Functions Transcending X-Chromosome Inactivation"

**A)** XACT expression in patients

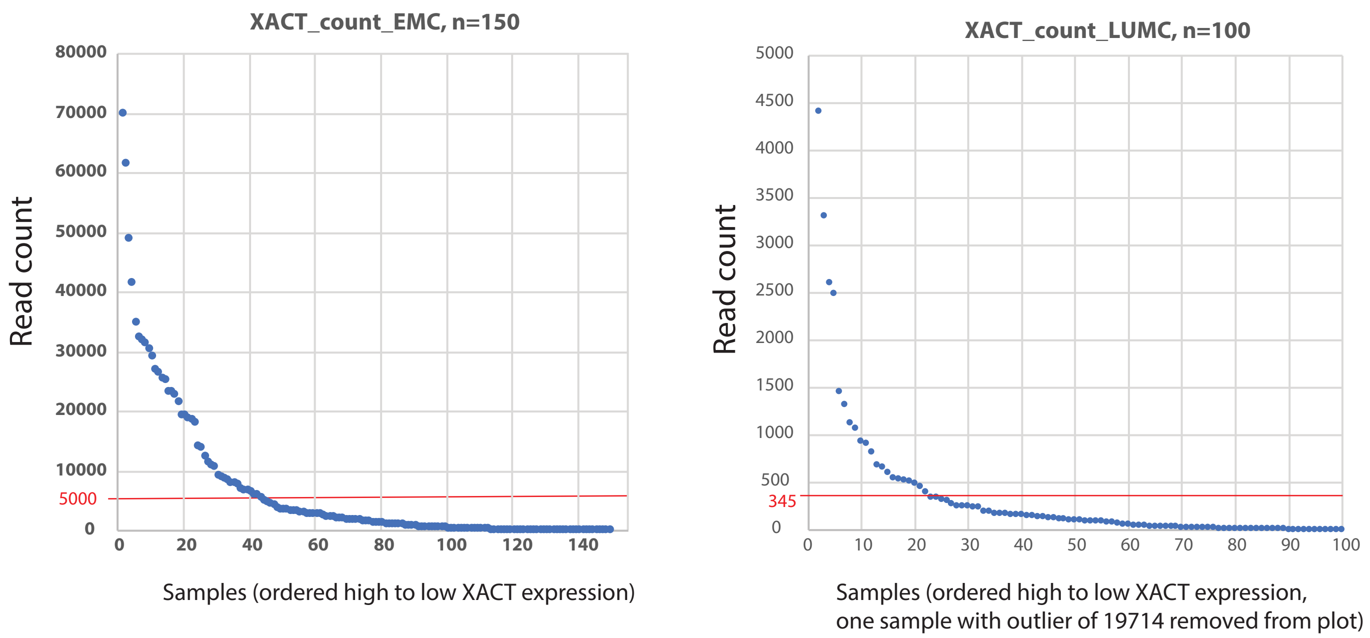


Distribution of XACT count from high to low, used for deciding the cutoff for expression.

**B)** XACT expression in selected patients (RNA sequencing)

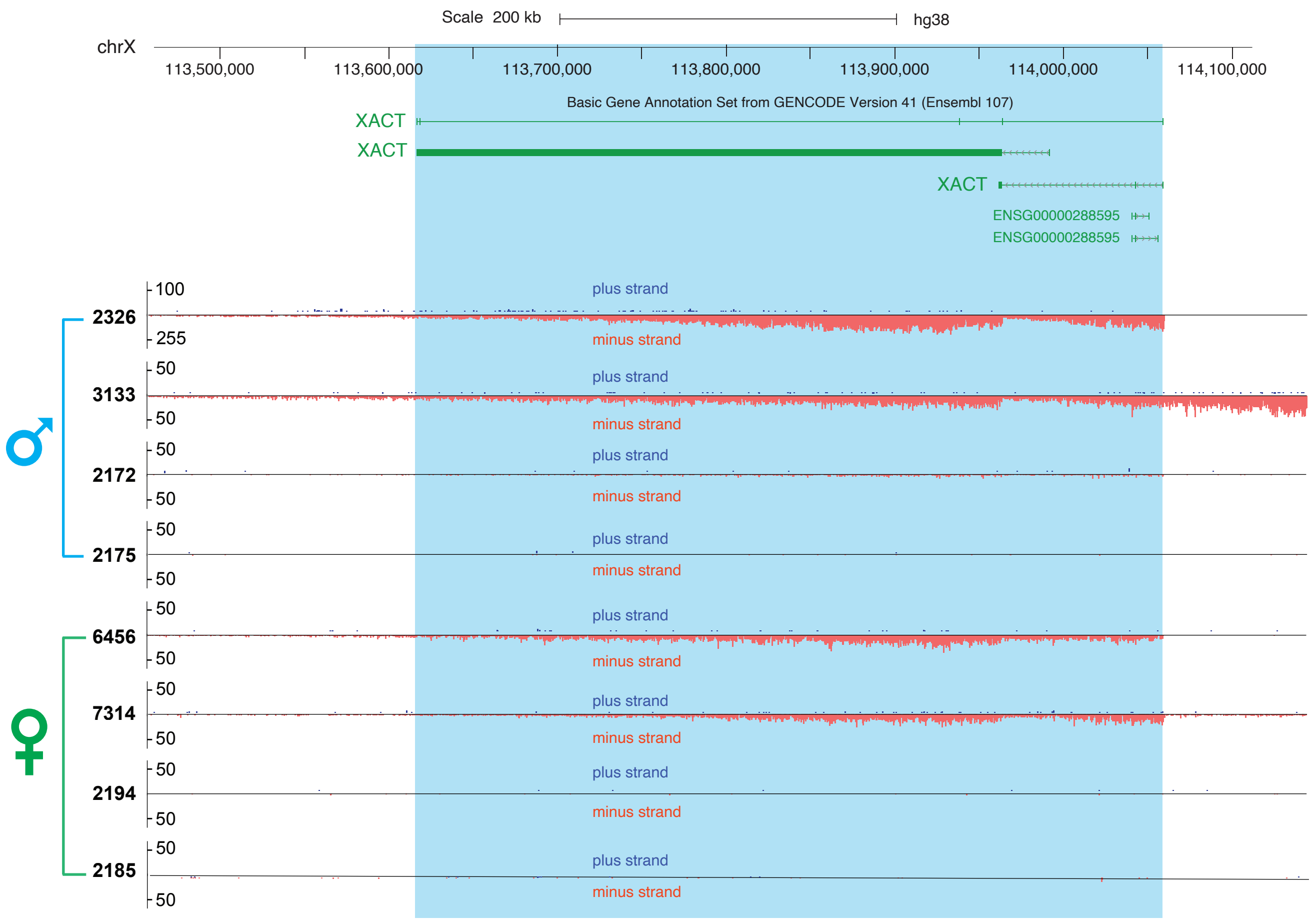


Zhang et al. 2025
