## Supplementary Figure_5 for "A Novel *XACT* lncRNA Transcript with Functions Transcending X-Chromosome Inactivation"

### Chromosome 3: LUMC dataset

**A)** Average expression of X linked genes across XACT high and low male patients

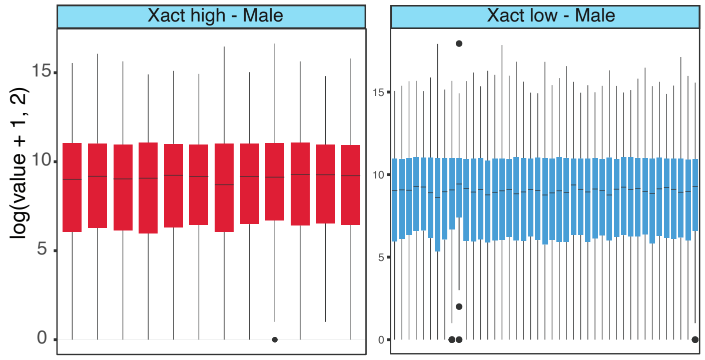


**B)** Average expression of X linked genes across XACT high and low female patients

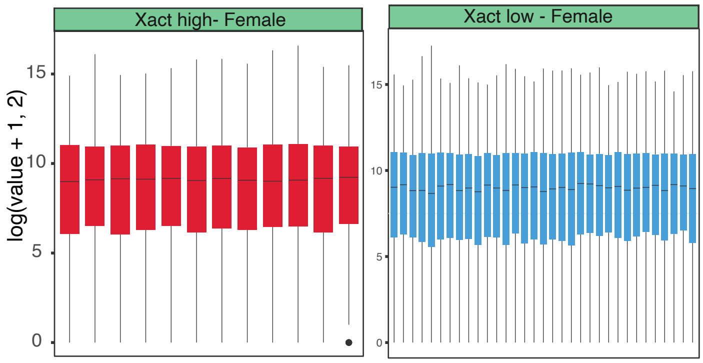


### Chromosome 8: LUMC dataset

**C)** Average expression of X linked genes across XACT high and low male patients

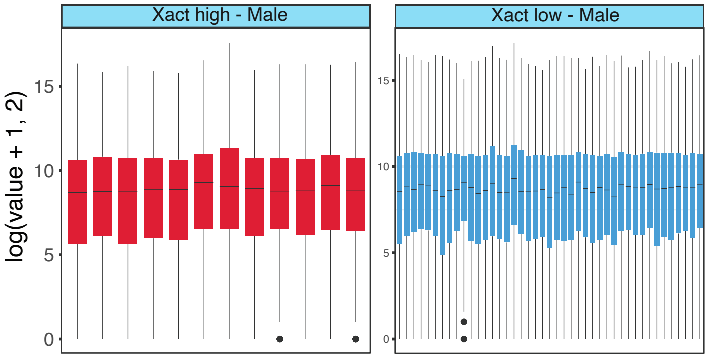


**D)** Average expression of X linked genes across XACT high and low female patients

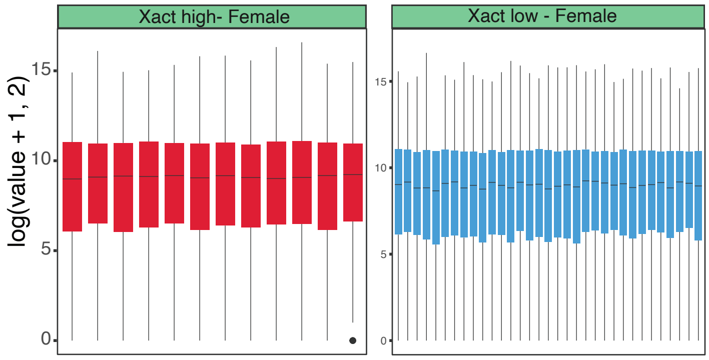
