## Supplementary Figure_6 for "A Novel *XACT* lncRNA Transcript with Functions Transcending X-Chromosome Inactivation"

**A)** XACT expression in different hematopoietic cells

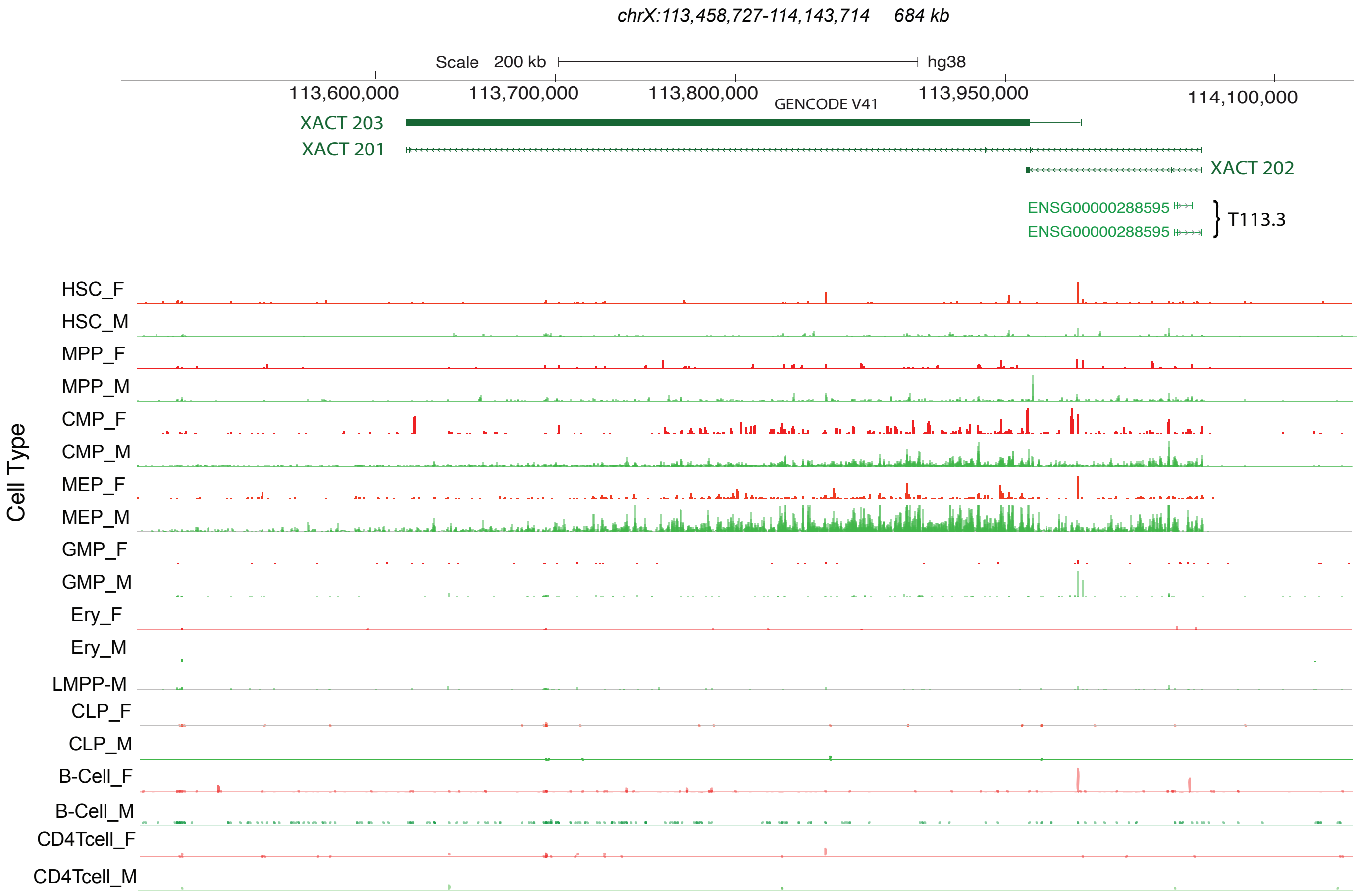


**B)** The chromatin dynamics of XACT locus

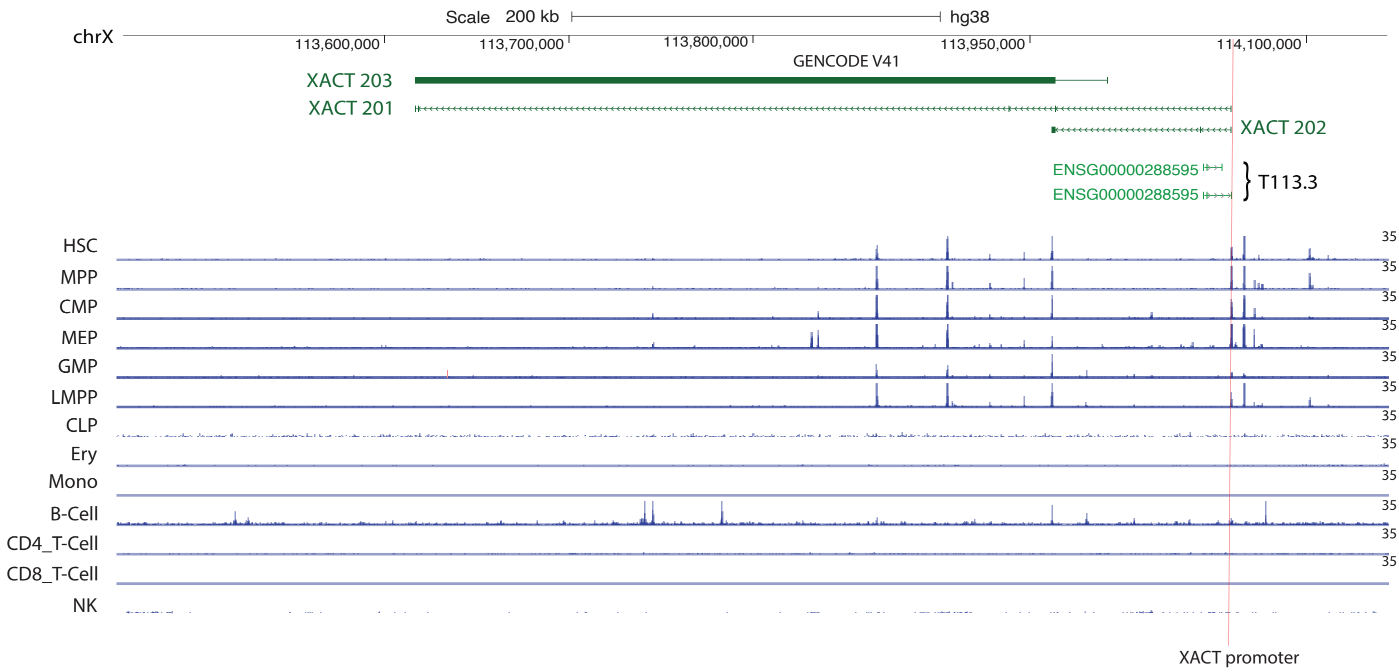


Zhang et al. 2025
