## Supplementary Figure_7 for "A Novel *XACT* lncRNA Transcript with Functions Transcending X-Chromosome Inactivation"

**A)** X chromosome to Autosome ratio (X:A ratio) in different hematopoietic cells in male and female donors

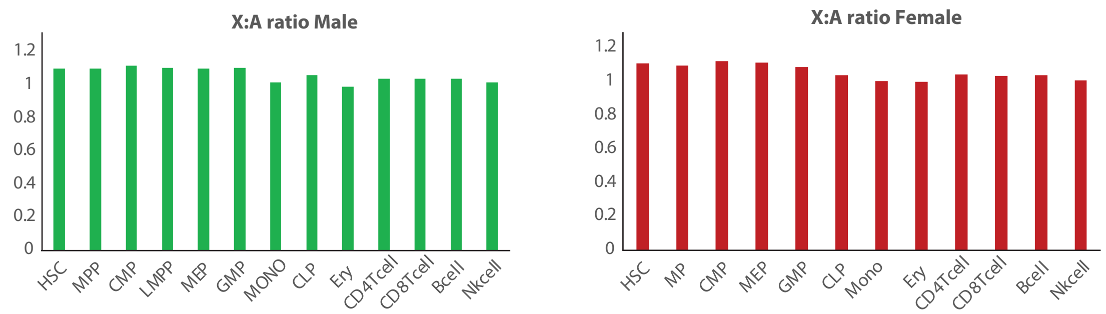


**B)** XACT expression in different hematopoietic cells
Dataset GSE113182

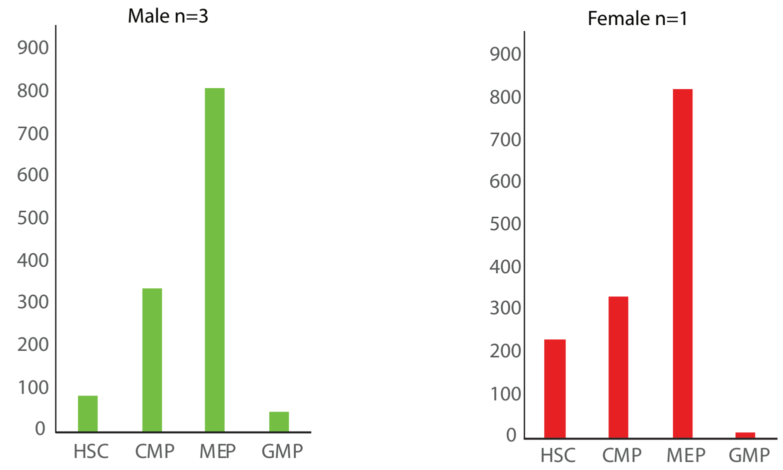


Zhang et al. 2025
