## Supplementary_Note_1.pdf for "A Novel *XACT* lncRNA Transcript with Functions Transcending X-Chromosome Inactivation"

**XACT is important for the recruitment of *cis*-regulatory element binding proteins and for transcriptional regulation.**

To gain a more detailed understanding of the role of XACT, we followed a multi-omics approach investigating chromatin accessibility (using ATAC-seq), changes in H3K27ac and H3K27me3 histone profiles (using CUT&Tag), and RNA-seq, and compared HUDEP2-KD to HUDEP2-NT cells.

First, in the chromatin accessibility data, when we analyzed and compared the number of differentially accessible regions (DARs) (with |log2FC| >= 1 and FDR <= 0.05), we obtained 685 regions (peaks) with higher accessibility in HUDEP2-KD (compared to HUDEP2-NT), and 741 regions with lower accessibility (Figure 5A; Supplementary Figure 8D; Supplementary Table 5). These DARs located across the genome, such as at promoters, introns, and distal intergenic regions. The highly enriched regions in HUDEP2-KD cells exhibited fewer peaks in the promoter within 1kb (Figure 5B), and they did not show a preference for the X-chr (Supplementary Figure 8G).

Next, we performed TF-occupancy (footprint) prediction using TOBIAS [1], an assay that uses the local distribution of Tn5 insertions in chromatin accessibility data to infer information about TF binding (caused by the visible depletion of insertions around sites bound by protein [1]). We identified 39 TFs with footprints that were more occupied in the HUDEP2-KD cells, and 43 TFs with higher occupancy in the HUDEP2-NT cells (fold change >95% or < 5% quantile, or -log10(p-value) > 95% quantile) (Figure 5C; Supplementary Table 5). The TF footprints that showed significant lower binding in HUDEP2-KD (compared to HUDEP2-NT) include TFs of the ETS family (e.g. SPI1, SPIB, FLI1, ETV1, ETV4) and other TFs such as TAL1, CEBPA, CEBPB, and CEBPD (Figure 5C; Supplementary Table 5). *SPI1* and *CEBPA* are both among the down-regulated genes in HUDEP2-KD cells, and SPI1 is the hub in the protein network analysis (shown in Figure 4H). SPI1 and the CEBP family proteins have been shown to be important for the proliferation as well as differentiation of erythrocyte progenitors [2]. SPI1 is essential for myeloid lineage commitment, functions as a suppressor of erythroid differentiation, and absence of SPI1 can lead to defective differentiation [3-7]. CEBPA has been shown to regulate erythroid differentiation negatively in MEL cells [2].

Inversely, TFs that showed significantly higher binding in HUDEP2-KD cells include the activator protein-1 (AP-1) family proteins (e.g., JUN, JUNB, JUND, ATF3, BATF, FOSB, FOSL2, and FOS) (Figure 5C; Supplementary Table 5). AP-1 family proteins play an important role in erythroid differentiation by activating erythroid-specific genes, and do promote erythropoiesis [8] [9-13]. This is consistent with the protein network analysis results wherein we found JUN as a hub in among the up-regulated genes in HUDEP2-KD cells (Figure 4G). We complemented this TF footprint analysis with motif analysis of the regions under DARs, and obtained enrichment of the ETS family members (such as SPI1, SPIB, FLI1) in DARs that show lower accessibility in HUDEP2-KD, and motifs of AP-1 family in DARs that show higher accessibility in HUDEP2-KD (Supplementary Table 5).

To further study the impact of *XACT* expression on chromatin, we performed Cut&Tag [62] with antibodies against H3K27ac (for active regions; enhancers and promoters) and H3K27me3 (inactive regions) on the aforementioned HUDEP2-NT and HUDEP2-KD cells. For H3K27ac, we identified 197 regions (peaks) that show significant changes in H3K27ac (|log2FC| >= 0.5, FDR <= 0.05) between HUDEP2-KD and HUDEP2-NT (84 with high H3K27ac in HUDEP2-KD, and 113 with low H3K27ac) (Figure 5D; Supplementary Figure 8E; Supplementary Table 6). The sites with low H3K27ac changes in the HUDEP2-KD group were frequently distributed in the promoter(<=1kb) region (Figure 5E, part *Down in HUDEP2-KD*, and their occurrence is more than two-fold increase, compared to those with high H3K27ac changes in the HUDEP2-KD group (Figure 5E, part *Up in HUDEP2-KD*). This further points to functions of XACT in recruiting TFs and transcriptional regulation. Similar to the ATAC-seq data, these regions display no specific preference for one chromosome (Supplementary Figure 8G).

Although the footprint analysis is optimized for chromatin accessibility data, we also applied it on H3K27ac Cut&Tag data, and obtained a TF footprint profile that is similar to that of chromatin accessibility (Figure 5F). TFs that show significantly higher binding in HUDEP2-KD, H3K27ac data include the AP-1 family TFs (e.g., JUN, JUNB, JUND, ATF3, BATF, FOSB, FOSL2, and FOS) and other TFs such as GATA1, GATA2 (Figure 5F; Supplementary Table 6). TFs that show significantly lower binding in HUDEP2-KD H3K27ac data include several ETS family TFs (e.g. ELK1, ETV1, GABPA, ELF1, EHF, ELF2, SPI1), and also MYB, and SP2. When we did a motif search on these H3K27ac-marked differential binding regions, we obtained similar results to the footprint analysis of the H3K27ac data and chromatin accessibility data. In the 84 regions with increased H3K27ac, we identified among others motifs for the AP1 family of TFs (e.g. JUN, JUNB, ATF3, BATF, FOSB, FOSL2, and FOS) (Supplementary Table 6). The 113 regions that show decreased H3K27ac contain motifs of the ETS family ( e.g. GABPA, ETV4, FLI1, ELK1, ETV1), and other TFs such as KLF1, MYB, SP1, SP2 (Supplementary Table 6).

In the H3K27me3 data, we identified 1243 regions with significant H3K27me3 modification (|log2FC| >= 1 and FDR <= 0.05), with 581 regions showing higher H3K27me3 in HUDEP2-KD compared to HUDEP2-NT, and 662 regions showing lower H3K27me3 (Figure 5G; Supplementary Figure 8F; Supplementary Table 7). These differentially H3K27me3 binding regions are distributed throughout the genome without chromosome preference (Figure 5H). Similar to chromatin accessibility and H3K27ac, these regions with significant H3K27me3 variability are with no specific enrichment on the X-chr (Supplementary Figure 8G). Although the H3K27me3 peaks are broad, not suitable for footprint analysis and direct comparison with the chromatin accessibility data and H3K27ac data, the motif analysis showed a similar trend in the -/+ 200 bp regions around the TSS of genes (where their TSS are covered by the differential H3K27me3 binding regions; Supplementary Table 7). We obtained PU.1 (SPI1), a TF belonging to the ETS family, in the 193 peaks around TSS that show higher H3K27me3 in HUDEP2-KD. We also found motifs for AP-1 family members (e.g. JUNB, BATF) in the 229 peaks around TSS that show lower H3K27me3 (Supplementary Table 7) in HUDEP2-KD compared to HUDEP2-NT.

To gain a more comprehensive understanding of XACT's function(s), we performed pathway enrichment analysis using a large list of gene sets obtained from RNA-seq, as well as the genes annotated using chromatin accessibility and histone modification results. Based on the HUDEP2-KD to HUDEP2-NT comparison, we grouped genes in two groups as follows: Group 1: downregulated (RNA-seq differential expression analysis), low chromatin accessibility (ATAC-seq), low H3K27ac, and high H3K27me3; and, inversely, Group 2: upregulated genes/RNAs, high chromatin accessibility, high H3K27ac, and low H3K27me3. For Group 1, the most significantly enriched pathways encompassed DNA-binding and TF activity (Figure 5I; for more details, Supplementary Figure 9; Supplementary Table 8), including *cis*-regulatory region binding, Transcription by RNAPol-II, DNA-binding, and DNA-templated transcription. This finding enhances the possible function(s) of XACT in recruiting TFs and its role in regulation of target gene transcription. We did not observe such enrichment in Group 2. Rather the pathways mainly related to response to stimulus and anatomical structure morphogenesis (Supplementary Figure 10; Supplementary Table 8).

In conclusion, our multi-omics approach shows that the function(s) of XACT extend beyond its effects on the X-chr, and affects several regions throughout the genome. The absence of XACT leads to changes in the binding of several TFs important for the proliferation as well as differentiation of erythroid progenitors. Specifically, XACT is proposed to recruit TFs to *cis*-regulatory regions and acts in transcriptional regulation.


1 Bentsen, M. *et al.* ATAC-seq footprinting unravels kinetics of transcription factor binding during zygotic genome activation. *Nat Commun* **11**, 4267 (2020). https://doi.org:10.1038/s41467-020-18035-1

2 Suh, H. C. *et al.* C/EBPalpha determines hematopoietic cell fate in multipotential progenitor cells by inhibiting erythroid differentiation and inducing myeloid differentiation. *Blood* **107**, 4308-4316 (2006). https://doi.org:10.1182/blood-2005-06-2216

3 Burda, P., Laslo, P. & Stopka, T. The role of PU.1 and GATA-1 transcription factors during normal and leukemogenic hematopoiesis. *Leukemia* **24**, 1249-1257 (2010). https://doi.org:10.1038/leu.2010.104

4 Nerlov, C., Querfurth, E., Kulessa, H. & Graf, T. GATA-1 interacts with the myeloid PU.1 transcription factor and represses PU.1-dependent transcription. *Blood* **95**, 2543-2551 (2000).

5 Zhang, P. *et al.* Negative cross-talk between hematopoietic regulators: GATA proteins repress PU.1. *Proc Natl Acad Sci U S A* **96**, 8705-8710 (1999). https://doi.org:10.1073/pnas.96.15.8705

6 Back, J., Dierich, A., Bronn, C., Kastner, P. & Chan, S. PU.1 determines the self-renewal capacity of erythroid progenitor cells. *Blood* **103**, 3615-3623 (2004). https://doi.org:10.1182/blood-2003-11-4089

7 Kato, H. & Igarashi, K. To be red or white: lineage commitment and maintenance of the hematopoietic system by the "inner myeloid". *Haematologica* **104**, 1919-1927 (2019). https://doi.org:10.3324/haematol.2019.216861

8 Jacobs-Helber, S. M., Wickrema, A., Birrer, M. J. & Sawyer, S. T. AP1 regulation of proliferation and initiation of apoptosis in erythropoietin-dependent erythroid cells. *Mol Cell Biol* **18**, 3699-3707 (1998). https://doi.org:10.1128/MCB.18.7.3699

9 Rosson, D. & O'Brien, T. G. AP-1 activity affects the levels of induced erythroid and megakaryocytic differentiation of K562 cells. *Arch Biochem Biophys* **352**, 298-305 (1998). https://doi.org:10.1006/abbi.1998.0597

10 Wu, Z., Nicoll, M. & Ingham, R. J. AP-1 family transcription factors: a diverse family of proteins that regulate varied cellular activities in classical hodgkin lymphoma and ALK+ ALCL. *Exp Hematol Oncol* **10**, 4 (2021). https://doi.org:10.1186/s40164-020-00197-9

11 Dore, L. C. & Crispino, J. D. Transcription factor networks in erythroid cell and megakaryocyte development. *Blood* **118**, 231-239 (2011). https://doi.org:10.1182/blood-2011-04-285981

12 Sumiko Takao, V. M., Fiona C. Brown, Richard Koche, Alex Kentsis. The AP-1/ETS Transcription Factor Network Regulates Stem Cell Quiescence and Chemotherapy Resistance in Acute Myeloid Leukemia. *Blood* **140** https://doi.org:https://doi.org/10.1182/blood-2022-165325

13 Sumiko Takao, V. M., Masahiro Uni, Alicia Slavit, Sophia Rha, Shuyuan Cheng, Laura K Schmalbrock, Fiona C Brown, Sergi Beneyto-Calabuig, Richard P Koche, Lars Velten, Alex Kentsis. Epigenetic mechanisms controlling human leukemia stem cells and therapy resistance. (2024). https://doi.org:https://doi.org/10.1101/2022.09.22.509005
